# Fine-mapping the Favored Mutation in a Positive Selective Sweep

**DOI:** 10.1101/139055

**Authors:** Ali Akbari, Joseph J. Vitti, Arya Iranmehr, Mehrdad Bakhtiari, Pardis C. Sabeti, Siavash Mirarab, Vineet Bafna

**Affiliations:** Department of Electrical & Computer Engineering, University of California, San Diego, La Jolla, CA 92093, USA; Department of Organismic and Evolutionary Biology, Harvard University, Cambridge, MA 02138, USA; Broad Institute of MIT and Harvard, Cambridge, MA 02142, USA; Department of Computer Science & Engineering, University of California, San Diego, La Jolla, CA 92093, USA

## Abstract

Methods to identify signatures of selective sweeps in population genomics data have been actively developed, but mostly do not identify the specific mutation favored by the selective sweep. We present a method, iSAFE, that uses a statistic derived solely from population genetics signals to pinpoint the favored mutation even when the signature of selection extends to 5Mbp. iSAFE was tested extensively on simulated data and in human populations from the 1000 Genomes Project, at 22 loci with previously characterized selective sweeps. For 14 of the 22 loci, iSAFE ranked the previously characterized candidate mutation among the 13 highest scoring (out of ∼ 21, 000 variants). Three loci did not show a strong signal. For the remaining loci, iSAFE identified previously unreported mutations as being favored. In these regions, all of which involve pigmentation related genes, iSAFE identified identical selected mutations in multiple non-African populations suggesting an out-of-Africa onset of selection. The iSAFE software can be downloaded from https://github.com/alek0991/iSAFE.

## Introduction

Genetic data from diverse human populations have revealed a multitude of genomic regions believed to be evolving under positive selection. We consider a regime where a single, *favored*, mutation increases in frequency in response to a selective pressure. The favored mutation either exists as standing variation at the onset of selection pressure, or arises *de novo*, after the onset. Neutral mutations on the same lineage as the favored mutation, hitchhike (are co-inherited) with the favored mutation, and increase in frequency, leading to a loss of genetic diversity.

Methods for detecting genomic regions under selection from population genetic data exploit a variety of genomic signatures. Allele frequency based methods analyze the distortion in the site frequency spectrum; Linkage Disequilibrium (LD) based methods use extended homozygosity in haplotypes; population differentiation based methods use difference in allele frequency between populations; and finally, composite methods combine multiple test scores to improve the resolution^1,2^. Recently, a lack of rare (singleton) mutations has been used to detect very recent selection^3^. The signature of a selective sweep can be captured even when standing variation or multiple *denovo* mutations create a ‘soft’ sweep of distinct haplotypes carrying the favored mutation. Together with the advent of deep sequencing, these methods have identified multiple regions believed to be under selection in humans and other organisms, and provide a window into genetic adaptation and evolution.

In contrast, little work has been done to identify the favored mutation in a selective sweep. Grossman et al.^4^ note that different selection signals identify overlapping but different regions, and a composite of multiple signals (CMS) can localize the site of the favored mutation. An alternative strategy is to use functional information to annotate SNPs and rank them in order of their functional relevance. However, the signal of selection is often spread over a large region, up to 1–2 Mbp on either side ^5^, and the high LD makes it difficult to pinpoint the favored mutation. Here, we propose a method, iSAFE (integrated <u>S</u>election of <u>A</u>llele <u>F</u>avored by <u>E</u>volution), that exploits coalescent based signals in ‘shoulders’ ^5^ of the selective sweep (genomic regions proximal to the region under selection, but carrying the selection signal) to rank all mutations within a large (5Mb) region based on their contribution to the selection signal. iSAFE requires that the broad region under selection is identified using existing methods, but does not depend on knowledge of the specific phenotype under selection, and does not rely on functional annotations of mutations, or knowledge of demography.

## Results

iSAFE uses a 2-step procedure to identify the favored variant, given a large region (5Mb) under selection. In the first step, it finds the best candidate mutations in small (low recombination) windows. Finally, it combines the evidence to give an iSAFE-score to all variants in the large region. It considers only biallelic sites, taking as input a binary SNP matrix with each row corresponding to a haplotype *h*, each column to a site *e*. Entries in the matrix correspond to the allelic state, with 0 denoting the ancestral allele, and 1 denoting the derived allele.

A haplotype ‘contains/carries a mutation *e*’ if it has the derived allele at site *e*. Recently, we devised the *Haplotype Allele Frequency* (HAF) score to capture the dynamics of a selective sweep^6^. The HAF score for a haplotype *h* (HAF(*h*)) is the sum of the derived allele counts of mutations in *h* (Fig. 1A and online methods). It has been shown that, when *h* is a carrier of the favored allele, HAF(*h*) increases with the frequency of the favored mutation (Eq. S9), in contrast to HAF scores of non-carriers (Eq. S10), and this can be used to separate carrier haplotypes from non-carriers without knowing the favored mutation^6^.

**Figure 1:**
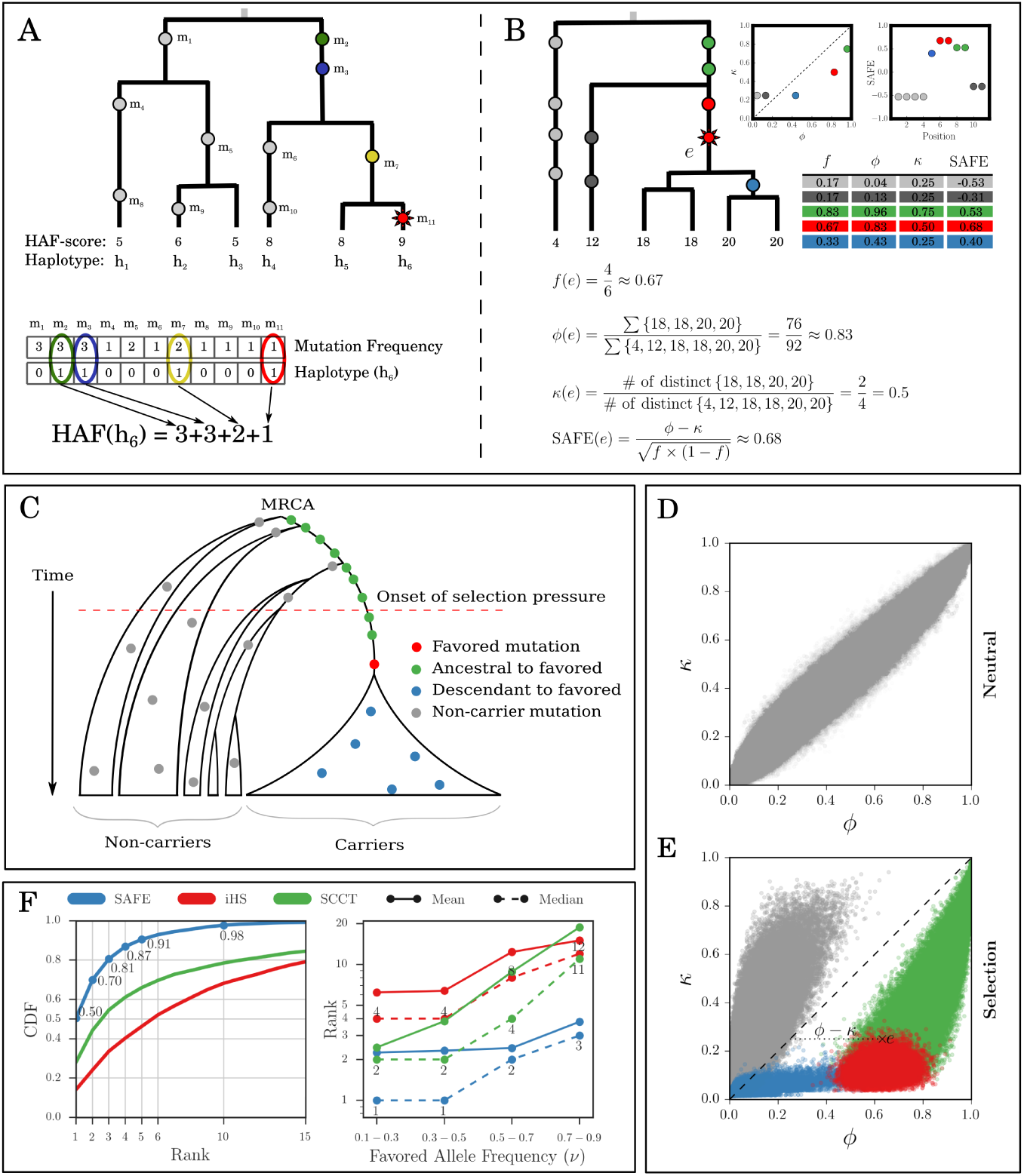
Illustration and Performance of the SAFE method. (**A**) The HAF score for haplotype *h* is the sum of the derived allele counts of the mutations on *h*. (**B**) Carriers of the favored mutation have higher fraction of the total HAF score of the sample (high *ϕ*) ^6^, and lower number of distinct haplotypes compared to non-carriers (low *κ*). (**C**) Schematic of a no-recombination (for exposition purposes) genealogy under a selective sweep. The mutations can be categorized as ‘non-carrier’ (gray), ‘ancestral to favored’ (green) arising prior to the favored mutation, and ‘descendant to favored’ (blue) that arise on haplotypes carrying the favored mutations but after the favored mutation, and the favored mutation itself (red). (**D, E**) Simulations showing *ϕ* versus *κ* values for each variant in a neutral evolution and a selective sweep with default parameters. The joint-distribution of *ϕ* and *κ*, in a selective sweep, changes in a dramatic but predictable manner that separates out non-carrier (gray), descendant (blue), and ancestral (green) mutations from the favored (red) mutations. The SAFE score computes a normalized difference of the two statistics. (**F**) Performance (favored mutation rank) of SAFE compared to iHS and SCCT on 50kbp windows with 1000 simulations per frequency bin. All the parameter have the default values for a fixed population size with ongoing selective sweeps. The left panel combines all allele frequencies while the right panel shows median and mean ranks for replicates divided into four bins.

Denote two haplotypes as ‘distinct’ if they have different HAF-scores. For any mutation *e*, let *f*_*e*_ denote the mutation frequency, or the fraction of haplotypes carrying the mutation. Let *κ*(*e*) (Fig. 1B) denote the fraction of distinct haplotypes that carry mutation *e*.

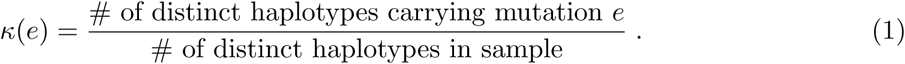

Similarly, let *ϕ*(*e*) denote the normalized sum of HAF-scores of all haplotypes carrying the mutation *e*.

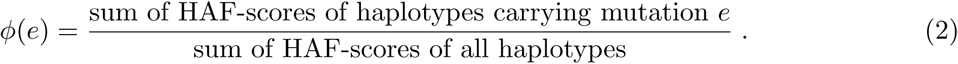

We observe empirically that in a region evolving according to a neutral Wright-Fisher model, *κ*(*e*) and *ϕ* (*e*) are both estimators of *f*_*e*_. Moreover, empirical results suggest that the expected value of *ϕ* (*e*) *-κ*(*e*) is 0, and variance is proportional to *f*_*e*_(1 *-f*_*e*_). Based on these observations, we define the SAFE-score of mutation *e* as

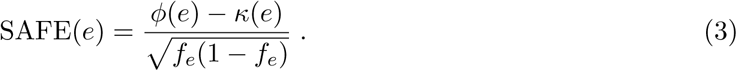

Empirically, SAFE(*e*) behaves like a Gaussian random variable, with mean 0, under neutrality (Fig. S2), and it can be used to test departure from neutrality. However, its real power appears during positive selection, when SAFE-scores change in a dramatic, but predictable manner (Fig. 1B-E). Assuming a no recombination scenario (only for visual exposition), label mutations as ‘non-carrier’ if they are carried only by haplotypes not carrying the favored allele. The remaining mutations can be labeled as ‘ancestral’, if they arise before the favored mutation, or ‘descendant’, if they arise after (Fig. 1C). Representing each mutation as a point in a 2-dimensional plot of *ϕ, κ* values, these classes are clustered differentially (Fig. 1D,E). The selective sweep reduces the number of distinct haplotypes carrying the favored mutation (lower *κ*), leaving non-carrier mutations with an increased fraction of distinct haplotypes (higher *κ*). On the other hand, increased HAF-scores in carrier haplotypes reduces the proportion of total HAF-score contributed by non-carrier haplotypes (lower *ϕ*). In contrast, the favored mutation has high positive value of *ϕ - κ* due to high HAF-scores for carriers (higher *ϕ*), and the reduced number of distinct haplotypes among its descendants (lower *κ*). As we go up to ancestral mutations, the number of non-carrier haplotype descendants increase, and *κ* grows faster than *ϕ*. As we go down to descendant mutations, there is a reduction in the already small number of distinct haplotypes. However, *ϕ* decreases sharply, reducing *ϕ - κ* (see Fig. 1B,C,E). Thus, we expect that the mutation with the highest SAFE-score is a strong candidate for the favored mutation.

We performed extensive simulations to test SAFE on samples evolving neutrally and under positive selection. We varied one parameter in each run (see online methods, ‘Simulation Experiments’), including window size (*L* = 50kbp), number of individual haplotypes (*n* = 200) chosen from a larger effective population size (*N* = 20*K*), scaled selection coefficient (*Ns* = 500), initial and final favored mutation frequencies (*v*_0_ = 1/*N*, and *v*). Only a few tests have been developed to identify or localize the favored mutation: Composite of Multiple Signals (CMS)^4^, and Selection detection by Conditional Coalescent Tree (SCCT)^7^. CMS combines statistics from different selection tests, including the integrated Haplotype Score (iHS)^8^, so as to localize the signal. In order to develop a unified probabilistic model, CMS expects control populations as input, as well as demographic models, and cannot be used directly on data based solely on coalescent simulations. Therefore, we compared SAFE against iHS and SCCT to obtain a baseline comparison here. The median SAFE rank of the favored mutation in a 50kbp region was 1 out of *∼*250 variants (left panel of Fig. 1F), and the favored mutation was in the top 5 in 91% of simulations. In comparison,the median ranks of iHS and SCCT were 6 and 3, respectively. Although SCCT was better at pinpointing selective causal sites in a small window (50kbp) than iHS, in larger regions (5Mbp), iHS performed better than SCCT (Fig. S13). The comparisons to CMS using simulated models of human demography are described later.

While standing variation, *v*_0_ > 1/*N*, generally weakens the selection signal, the performance of SAFE remains relatively robust to variation in *v*_0_. The median SAFE rank of the favored allele is at most 3 out of *∼*250 variants in all cases except when *v*_0_ *≥* 1000/*N* (Figure S4). Similarly, the performance is robust to selection pressure, with only a slight degradation at weak selection (*Ns* = 50) (Fig. S5) where the median rank goes to 9 (3.5%-ile), while for *Ns ≥* 200 the median rank is at most 2. As expected, the performance improves with increasing sample size (Fig. S6). We also tested SAFE on a model of European demography and observed similar results (Fig. S7). These tests used L = 50kbp, chosen so as to minimize the effects of recombination.

Next, we tested SAFE with increasing window sizes, and observed that while the median rank of the favored mutation increases with increasing window size, the percentile rank improves up to 80kbp and then degrades to 3%-ile around 1Mbp (Fig. 2A, and S8). The deterioration for larger windows is likely due to most haplotypes becoming unique, and *κ* losing its utility in pinpointing the favored mutation. However, the selective sweep signal is known to extend to large, linked regions, as far as 1Mbp on either side of the favored allele. These ‘shoulders’ of selective sweeps are helpful in identifying the region under selection, but make it harder to pinpoint the favored mutation. We further refined our method to exploit the signal from shoulders.

**Figure 2:**
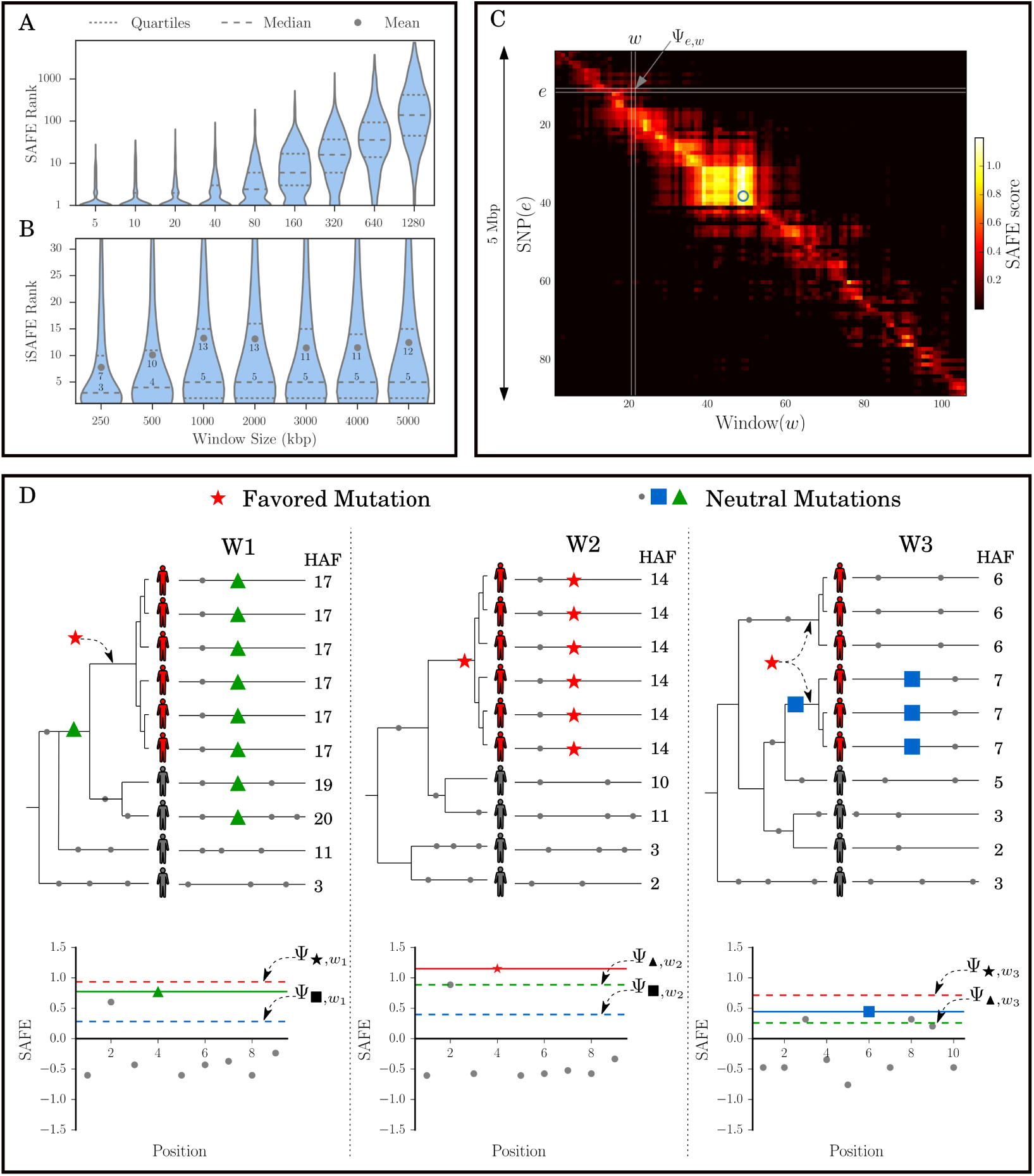
iSAFE. (**A**) SAFE performance (rank distribution of favored mutation) as a function of window-size. The dashed line represents median rank, and decays for large windows while (**B**) iSAFE is robust to increase in window size. (**C**) The Ψ(*e, w*) matrix, with *e ϵ S*1, for a 5Mbp region around LCT gene in FIN population shows that the ‘shoulder’ of selection can extend for a few Mbp. The blue circle shows the location of the putative favored mutation rs4988235. (**D**) The red-star, green-triangle, and blue-square denote the favored, ancestral, and descendant mutations, respectively. The order of haplotypes is preserved in the three windows. In its own window *w*1, the triangle has a small number of distinct haplotypes. However, when inserted into *w*2 or *w*3, the number of distinct haplotypes increases (increased *κ*). Analogously, when the square is inserted in *w*1 or *w*2, it reduces the proportion of total HAF-score (lower *ϕ*). In contrast, the star (favored) mutation, retains high *ϕ* and low *κ* when inserted in any windows.

For larger regions, we considered a set of 50% overlapping windows of fixed size (300 SNPs). For each window, we applied SAFE and chose the mutation with the highest SAFE-score. Let *S*_1_ denote the set of selected mutations. Mutations in *S*_1_ are likely to contain either the favored mutation itself or mutations linked to it. For mutation *e* in window *w*, let Ψ_*e,w*_*′* denote the larger of the SAFE-score of *e*, when *e* is ‘inserted’ into window *w*′, and 0 (Fig. 2C). As different windows have different genealogies due to recombination, Ψ_*e,w*_*′* is relatively high when *e* is the favored mutation and the genealogies of *w, w*′ are identical or very similar, but not otherwise. In contrast, the SAFE-score of a non-favored mutation *e* is relatively low when inserted in other windows (Fig. 2D; see online methods). Define the weight of a window *w* as

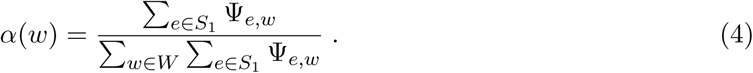

Windows that contain the favored mutation and those sharing its genealogy are expected to have high *a* values. We defined the iSAFE-score for all mutations *e* (including those not in *S*_1_) as:

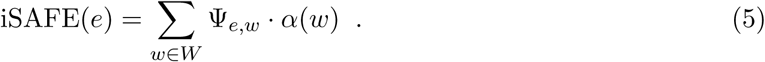

We tested the power of iSAFE to identify the favored mutation in varying window sizes and observed consistently high performance as the window size was increased from 250kbp all the way to 5Mbp (Fig. 2B). The median rank remained between 3 and 5 up to 5Mbp, and the performance remained robust to a large range of parameter choices including both hard and soft sweep scenarios, selection pressure and favored mutation locations (Fig. S11, S12). iSAFE greatly improved upon iHS and SCCT, placing the favored mutation within top 20 in 88% of the cases, in contrast to iHS (39%), and SCCT (34%), for an ongoing selective sweep with fixed population size (Fig. S13).

iSAFE-scores are not based upon likelihood computations, and the distribution of scores depend upon largely unknown factors including demography, time since onset of selection, selection coefficient, and other parameters. Nevertheless, they can be used to rank order the mutations. Additionally, iSAFE scores are normalized and can be compared across samples. We found distinct differences in performance below a score threshold of 0.1. The median rank of the favored mutation is 4 when peak iSAFE-score exceeds 0.1 versus a median rank of 10 along with a longer tail, when peak iSAFE-score is below 0.1 (Fig. S14). Empirically computed *p*-values (online methods) on iSAFE indicate good performance when *p*-value < 1e-4 (Fig. S15).

Not surprisingly, iSAFE performance deteriorates when the favored mutation is fixed, or near fixation (v > 0.9 in Fig. S16). To handle this special case, we include individuals from non-target populations. For a mutation, define the <u>M</u>aximum <u>D</u>ifference in <u>D</u>erived <u>A</u>llele <u>F</u>requency score (MDDAF) as the difference

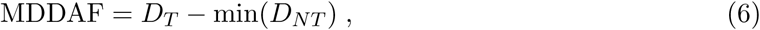

where *D*_*T*_ is the derived allele frequency in the target population and min(*D*_*NT*_) is the *minimum* derived allele frequency over all non-target populations. Simulations of human population demography under neutral evolution (Fig. S20), shows *P* (MDDAF > 0.78*|D*_*T*_ > 0.9) = 0.001 (see Fig. S22). Therefore, when we observe the rare event of high frequency mutations in target (*D*_*T*_ > 0.9) with MDDAF > 0.78, we add random outgroup samples to the data to constitute 10% of the data (online methods). In testing on the phase 3 of 1000 Genomes Project (1000GP) data, we chose outgroup samples from non-target 1000GP populations. The addition of outgroup samples using the MDDAF criterion was tested in extensive simulations. While the performance did not change for *v* < 0.9, it dramatically improved for high frequencies, including when the favored mutation was fixed in the target population (Fig. S16). In testing on models of human demography, we also compared against CMS. While CMS showed excellent performance in localizing the favored mutation, iSAFE scoring greatly improved the ranking. For example, iSAFE ranked the favored mutation within the top 20 in 94% of the simulations of a 5Mbp region (Fig. 3, S17), in contrast to CMS which had a top 20 ranking in 35% of cases.

**Figure 3:**
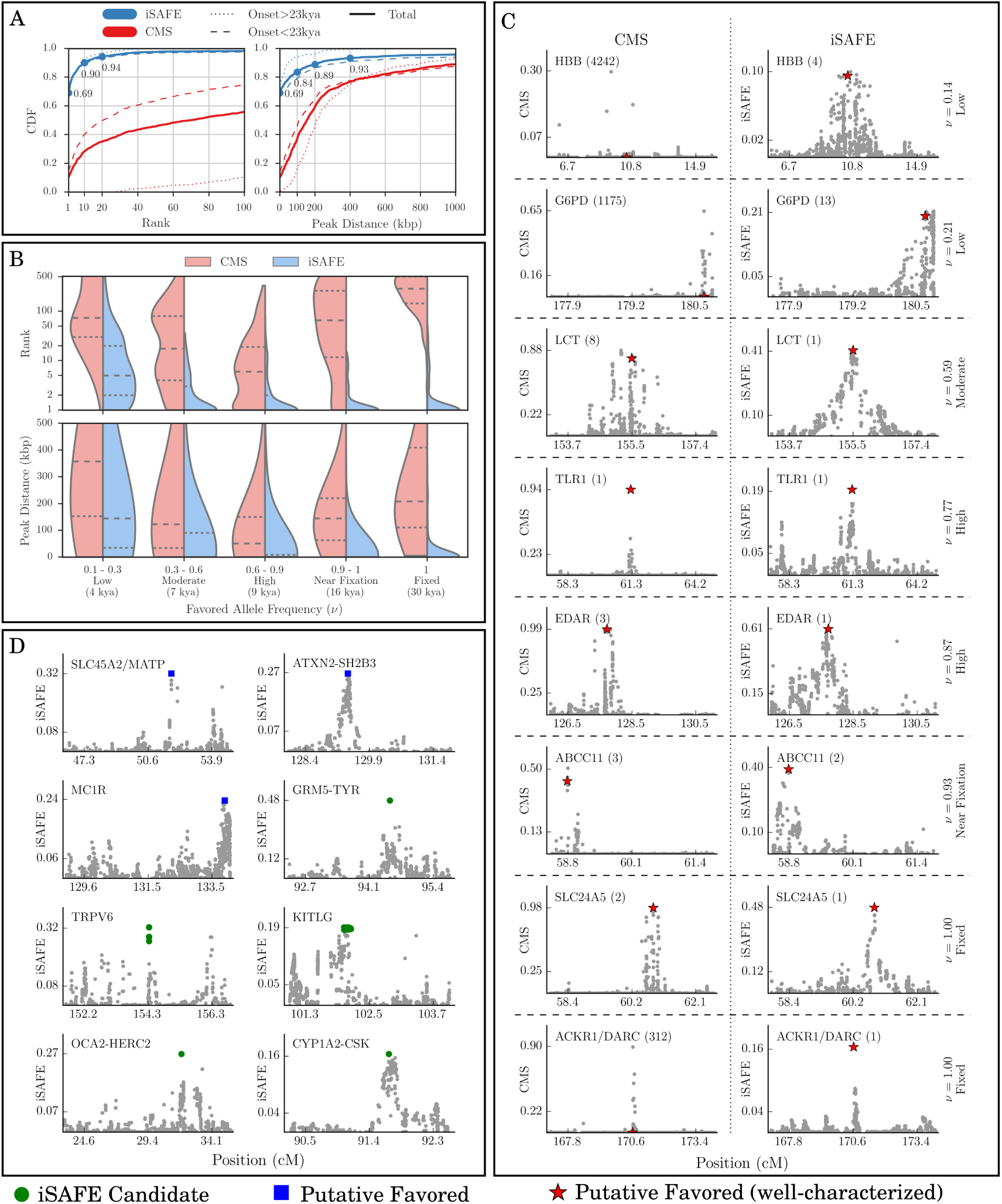
iSAFE Performance. (**A**) The left (right) panel is the Cumulative Distribution Function (CDF) of favored mutation rank (peak distance) for iSAFE and CMS scores, shown by blue and red, respectively. All data is based on simulations of 5Mbp genomic regions simulated using a model of human genome based on the human demography (Fig. S20). The time of onset of selection was chosen at random (using the distribution in Fig. S21) after the out of Africa event, in the lineage of EUR population (as the target population). When the onset of selection is before split of EUR and EAS (> 23kya), both (EUR and EAS) are under selection. (**B**) iSAFE performance (rank and peak distance distributions of favored mutation) as a function of favored allele frequency (*v*) in the target population (EUR). The dashed (dotted) line represents median (quartiles). (**C**) iSAFE and CMS scores (right and left panels, respectively) on 8 well-characterized selective sweeps (Table S1). The rank of the putative favored mutation (red star) in 5Mbp region is shown in parentheses. (**D**) iSAFE-scores on regions under selection. Top ranked iSAFE candidates are marked by blue squares when they match putative favored mutations, while green circles represent new favored mutations suggested by iSAFE. All data-sets were chosen by taking a 5Mbp window around the putative selected region, unless one side reached the telomere or centromere.

In testing instances of previously characterized sweeps in 1000GP data, we note that perfor-mance is difficult to characterize due to many complicating factors. Multiple sweeps could be occurring in response to different selection events, including background selection in the same region; or polygenic selection may also dilute the selection signal at any one locus. Moreover, the favored mutation is well-characterized in only a few instances. We looked for genes/regions that showed the signature of a selective sweep in one of the 1000GP sub-populations, and had additional evidence pointing to the favored mutation. We identified 22 genes with some evidence, but only 8 ‘well characterized’ cases with additional support for the favored mutation (Supp. Table S1).

We used iSAFE to rank all variants (*∼*21,000) in a 5Mbp region surrounding the gene. Among the 8 well characterized cases, (Fig. 3C), iSAFE ranked the candidate mutations as 1 in 5 cases: SLC24A5, LCT, EDAR, ACKR1, TLR1; and, it assigned ranks 2 (ABCC1), 4 (HBB), and 13 (G6PD) in others. In almost all cases, we observed high iSAFE-scores (*≥*0.1).

We checked to see if the other 14 regions under selection showed a strong iSAFE signal. In 3 of the 14 regions (FUT2, F12, ASPM; Fig. S30), we only observed weak signals, and did not make a prediction (peak iSAFE < 0.027), although we do see a strong iSAFE peak 1.3Mbp away from the ASPM gene (Fig. S30D). In other regions, iSAFE ranked the candidate mutations as 1 in the SLC45A2/MATP (CEU), MC1R (CHB+JPT), and ATXN2-SHB3 (GBR) genes (Fig. 3D), and 7, 8, and 12 in PSCA (YRI), ADH1B (CHB+JPT), and PCDH15 (CHB+JPT) genes, respectively. In each case, the iSAFE-scores were high with the exception of PSCA (peak iSAFE = 0.04, online methods).

The other 5 putative selected regions are interesting in that the top-ranked iSAFE mutations had high scores, but were distinct from the reported candidate mutations (Fig. 3D). Many of these genes are involved in pigmentation, *deterMining*, skin, eye, and hair color. For example, the Tyrosinase (TYR) gene, encoding an enzyme involved in the first step of melanin production, is considered to be under positive selection with a nonsynonymous mutation rs1042602 as a candidate favored variant^9^. A second intronic variant, rs10831496, in GRM5, 396kbp upstream of TYR, has been shown to have a strong association with skin color ^10^. In contrast, iSAFE ranks mutation rs672144 at the top. Interestingly, this variant was the top ranked mutation not only in CEU (iSAFE = 0.48, *p*-val*≪*1.3e-8), but also in EUR, EAS, AMR, and SAS (iSAFE >0.5, *p*-val*≪*1.3e-8;Fig. S23). The result is consistent with the signal of selection being observed in all populations except AFR. It may not have been previously reported because it is near fixation in all populations of 1000GP except for AFR (Fig. S23H). We plotted the haplotypes carrying rs672144 and found that two distinct haplotypes carry the mutation, both remaining high frequency, maintained across a large stretch of the region, suggestive of a soft sweep with standing variation (Fig 4). A similar analysis applied to genes TRPV6, KITLG, OCA-HERC2 (Fig. 3D), where in each case, the top iSAFE mutations were identical across all non-African populations (online methods), and supported an out-of-Africa onset of selection. In the one remaining gene (CYP1A2/CSK; Fig. 3D), the top ranked iSAFE mutation rs2470893 was previously found significant in a genome wide association study^11^, and was tightly linked to the candidate mutation. To summarize, iSAFE analysis ranked the candidate mutation among the top 13 in 14 of the 22 loci, did not show a strong signal in 3, and identified plausible alternatives in the remaining 5.

**Figure 4:**
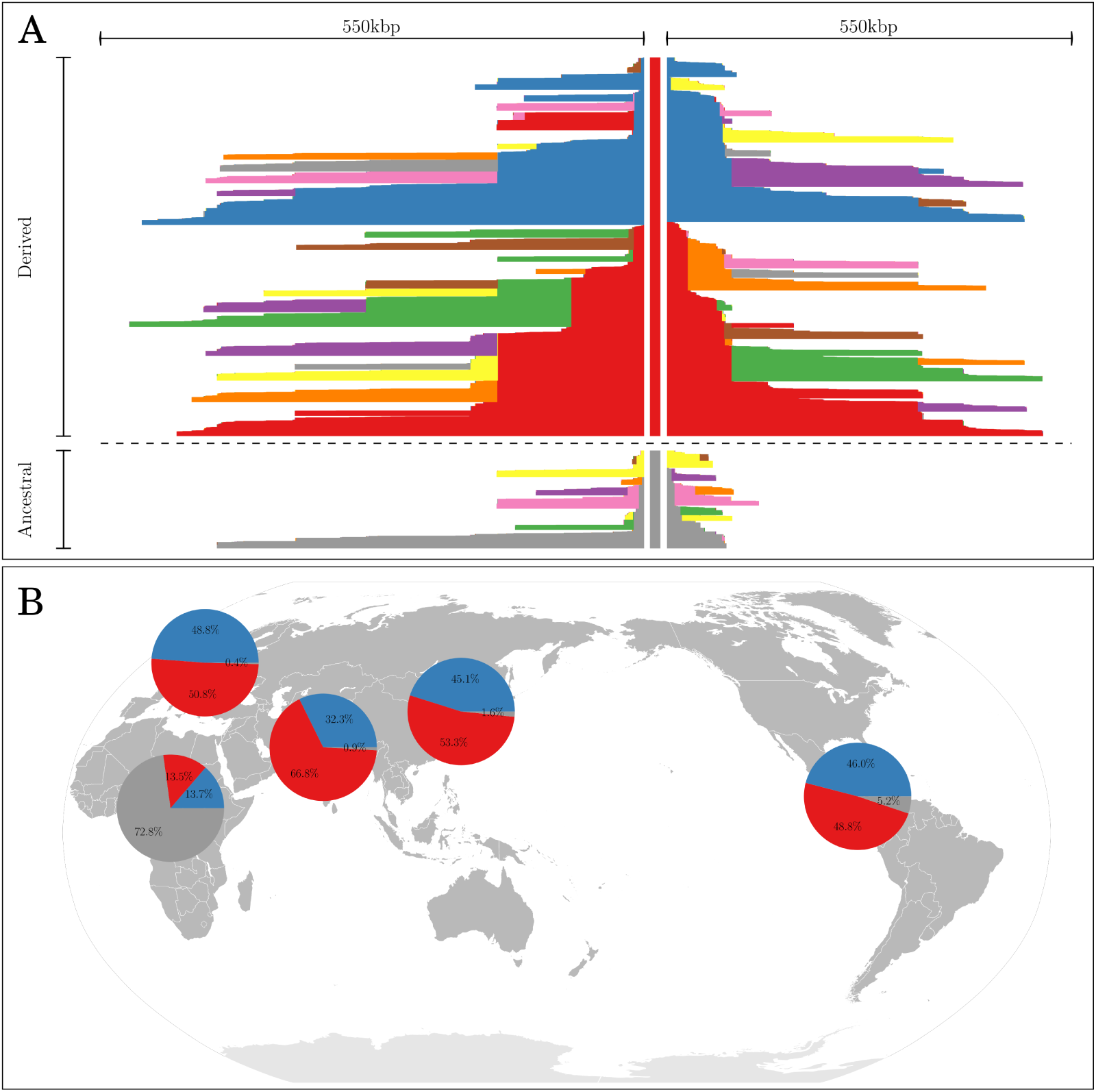
The GRM5-TYR haplotype structure. The mutation rs672144 is ranked first by iSAFE and very well separated from rest of the mutations in 5Mbp around it, in all non-African populations with high confidence(iSAFE >0.5, *p*-val*≪*1.3e-8). (**A**) Haplotype plot with core mutation rs672144 on all 2504 *×* 2 haplotypes of 1000GP. This plot shows carrier haplotypes of mutation rs672144 are conserved over a longer span than haplotypes in non-carriers which is a signal of selection ^8^. (**B**) Global frequencies of carrier haplotypes of mutation rs672144 (red, blue) and non-carrier haplotypes (gray). The evidence is consistent with an out of Africa selection on standing variation (soft sweep) with mutation rs672144 being the favored variant.

## Discussion

The identification of the favored allele in a selective sweep is a long-standing computational problem in population genomics. Our results suggest that statistics obtained from the coalescent structure of a region under a selective sweep can indeed pinpoint the favored mutation. iSAFE was designed to work in regimes where the selection strength is high, and there is a single favored mutation. However, its performance remains robust to a range of simulation parameters, including a wide range of initial frequencies (standing variation), and the frequency of the favored mutation at the time of sampling. iSAFE is not highly parametrized. While most results in the paper are presented on human populations, iSAFE can be easily extended to other populations, with additional demographic simulation or empirical calculations required to recalibrate p-values.

An important challenge was that regions undergoing a selective sweep also present a signal far away from the favored mutation, making it harder to pinpoint the favored mutation. We observe that when a true favored mutation is inserted into a shoulder region, it gets higher SAFE-scores on average, in contrast to the insertion of a hitchhiking mutation. The iSAFE technique uses this idea to exploit the shoulders and rank mutations according to the weighted sum of their SAFE-scores in all windows.

We also use a cross-population technique in a limited manner by using the frequency differential of mutations in high frequency scenarios to get representative non-carrier haplotypes in the sample, and show its power in identifying nearly fixed favored mutations. We do assume a model with a single, favored variant, and future work could contribute to identify multiple interacting loci favored by selection. Finally, we use only population based methods, and future work will seek to integrate these techniques with a functional analysis of mutations.

## Acknowledgments

This research was supported in part by grants from the NSF (IIS-1318386 and DBI-1458557), and from the NIH (R01GM114362).

## Online Methods

### 1 The iSAFE statistic

#### 1.1 iSAFE: Input, Output and Overview

Consider a sample of phased haplotypes in a genomic region. We assume that all sites are biallelic and polymorphic in the sample. Thus, our input is in the form of a binary SNP matrix with each row corresponding to a haplotype and each column to a mutation, and entries corresponding to the allelic state, with 0 denoting the ancestral allele, and 1 denoting the derived allele. The output is a non-negative iSAFE-score for each mutation, with the highest score corresponding to the favored mutation.

At a high level, iSAFE uses a 2-step procedure to identify the favored variant, given a large region (5Mb) under selection. In the first step, it finds the best candidate mutations in small (low recombination) windows. Finally, it combines the evidence to give an iSAFE-score to all variants in the large region.

#### 1.2 The <u>H</u>aplotype <u>A</u>llele <u>F</u>requency (HAF-)score

The HAF score for haplotype *h* is the sum of the derived allele counts of the mutations on *h*. Define the SNP matrix *M* such that, *M*_*h,e*_ = 1 if haplotype *h* carries the derived allele of SNP *e*, and 0 otherwise. The Haplotype Allele Frequency (HAF) score of haplotype *h* defined in Ronen et al. (2015)^6^ as:

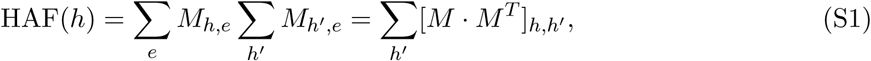

where 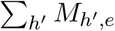 is derived allele count for SNP *e*, and 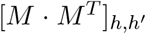 is number of shared derived alleles (mutations) between haplotypes *h* and *h*^*t*^ (see Fig. 1A).

#### 1.3 SAFE: **<u>S</u>election of <u>A</u>llele <u>F</u>avored by <u>E</u>volution.**

For each SNP *e*, define *ϕ* as:

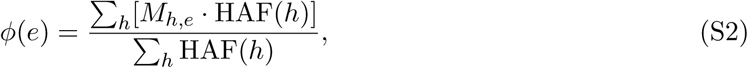

In other words, *ϕ* is sum of HAF scores of carriers of the derived allele *e* 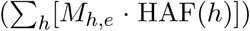, divided by sum of HAF scores of all haplotypes in the sample 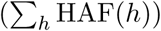.

Similarly, for each SNP *e*, we define *κ* as:

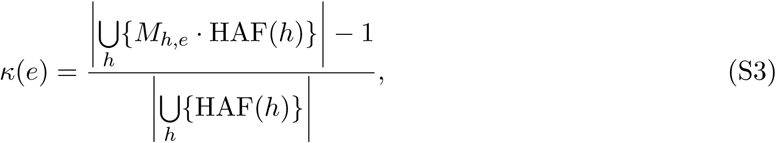

implying that *κ* is the fraction of distinct non-zero values in HAF scores of SNP *e* carriers. *κ* is closely related, but not identical, to fraction of all distinct haplotypes that carry the mutation *e*.

We use *ϕ* and *κ*, to define the SAFE score of a SNP *e* as:

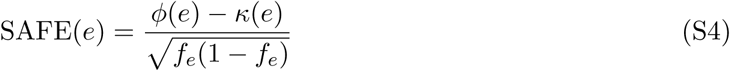

where *f*_*e*_ is the derived allele frequency of SNP *e*.

To explain the behavior of the SAFE-score in pin-pointing the favored mutation, we describe a collection of theoretical and empirical observations that can be summarized as follows:

1. *Under neutrality, ϕ*(*e*) *and κ*(*e*) *are (biased) estimators of f*_*e*_.
2. *λf* (1 – *f*) *is a biased estimator for variance of* (*ϕ – κ*), *where λ is a positive constant*.
3. *The two points above allow the use of SAFE-score as a statistic that empirically follows a Gaussian distribution with mean* 0 *under neutrality*.
4. *For a population evolving under selection, ϕ and κ move in opposite directions. Specifically, for the favored mutation e, ϕ*(*e*) *increases, while κ*(*e*) *decreases. The SAFE-score tends to be maximized for the favored mutation e*.

We elaborate on these points below.

##### 1.3.1 Behavior of *ϕ, κ* under neutrality, constant population size

Consider a sample of size *n* selected from a population evolving neutrally according to the Wright Fisher model (constant population size, random mating, discrete generations, no recombination), with scaled mutation rate *θ*. Let *ξ*_*i*_ be the number of sites with derived allele count *i*. From Ronen et al. ^6^, the mean of the HAF scores of all *n* haplotypes in the sample is

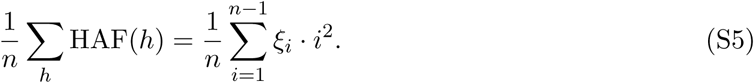

Under the coalescent model, Eq. (22) of Fu 1995^12^ shows that 𝔼[*ξ*_*i*_] = *θ/i* for all 1 *≤ i ≤ n* - 1. By averaging over all haplotypes in all genealogies, the expected HAF score is computed as

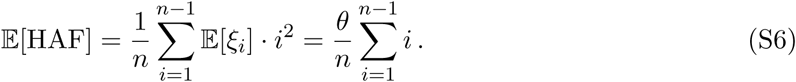

Thus, the expected HAF score is,

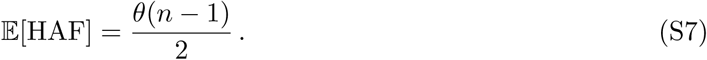

Therefore, the fraction of the total HAF-score of *fn* randomly chosen haplotypes is approximately *f*. A mutation *e* with derived allele frequency also has *fn* descendants (carriers). However, to compute the sum of the HAF-scores, we must consider a random coalescent process with a condition that carriers coalesce to a common ancestor before any carrier coalesces with a non-carrier.

**Figure S1:**
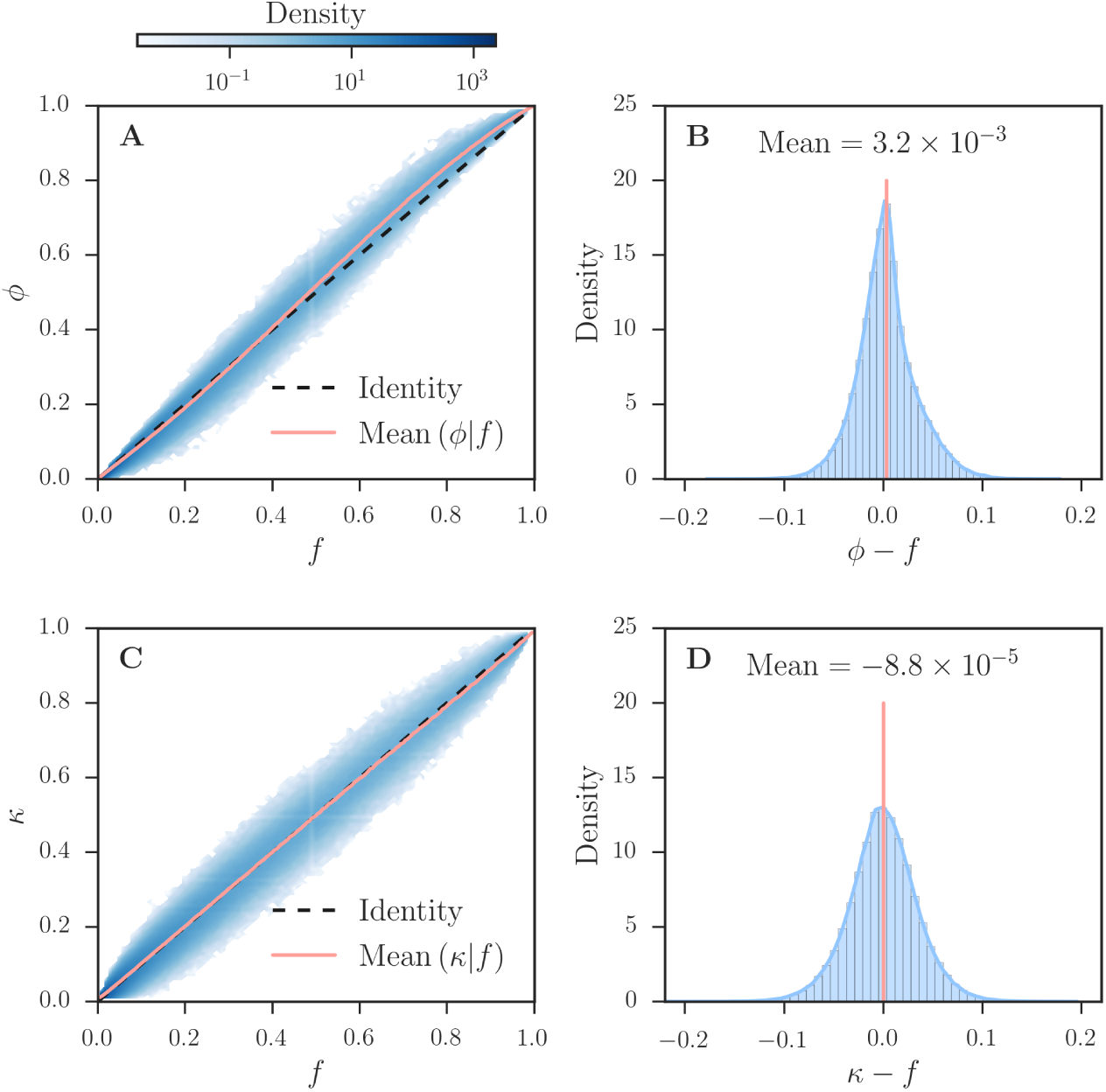
*κ* and *ϕ* as estimators of*f*. Empirical analysis, with 10,000 neutrally evolving population (about 3 million SNPs) with default parameter set, shows that *ϕ* and *κ* are (biased) estimators of allele frequency *f* (*f* = *i/n* for all integers *i ∈* [1, *n* - 1]).

This is harder, even though conditional coalescent processes have been studied extensively (e.g., Wiuf and Donnelly^13^). Empirical analysis on neutral coalescent simulations conditioned on the mutation *e* having *fn* carriers reveals that (Fig. S1)

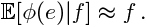

While *κ* has not been studied previously, it is closely related to the fraction of distinct haplotypes in the sample. Empirically, for a mutation *e*, with *fn* descendants, we observe that (Fig. S1)

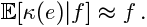

and, for all *e* (Fig. S2),

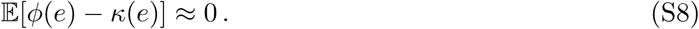

**Figure S2:**
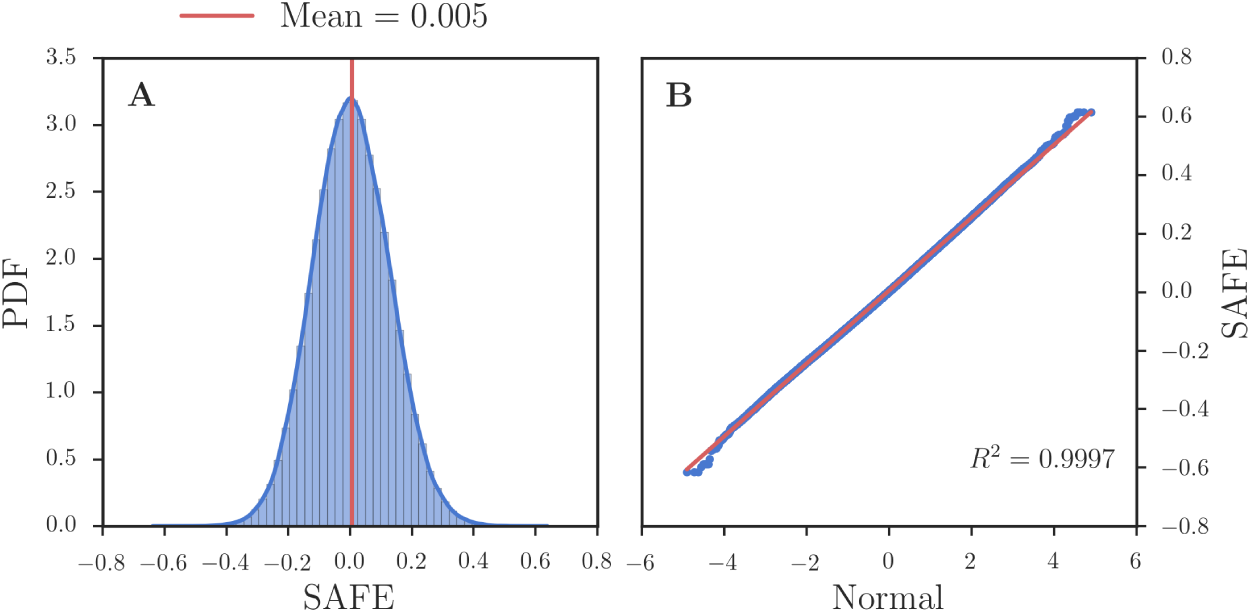
Empirical SAFE distribution. (**A**) SAFE score Probability Density Function (PDF) of 10,000 neutrally evolving population (about 3 million SNPs) with default parameter set. (**B**) Quantiles of the SAFE score against the quantiles of the Normal distribution for the same data in part A. The coefficient of determination (*R*^2^ = 0.9997) for the QQ-plot shows that Gaussian distribution is a good approximation to the SAFE score distribution.

##### 1.3.2 Distribution of SAFE-scores in a neutrally evolving population

The discussion above suggests that E(SAFE(*e*)) = 0 for all derived alleles *e*. Additionally, empirical observations suggest that *λf* (1*-f*) is a biased estimator for variance of (*ϕ-κ*), where *λ* is a positive constant. We observed empirically that the distribution of the SAFE score of derived alleles in a neutrally evolving population is therefore approximated by a *Gaussian* distribution with mean 0 and unknown variance *λ* (see Fig. S2).

##### 1.3.3 Behavior of *ϕ, κ*, SAFE in a population under selection, constant population size

The dynamics of HAF-score for a haplotype carrying the favored mutation in an ongoing selective sweep was analyzed earlier ^6^. It increases dynamically upto fixation of the favored allele, and then decreases dramatically.

Formally, let HAF^car^ (respectively, HAF^non^) denote the HAF score of a random haplotype carrier of the favored allele (respectively, a non-carrier) when a fraction *f* of the *n* sampled haplotypes carry the favored allele. In S1 Text of Ronen et al.(2015) ^6^, we show that under strong selection (*Ns "* 1) and no recombination,

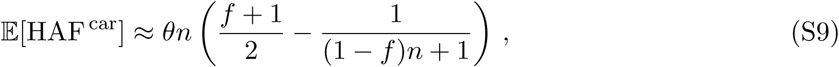

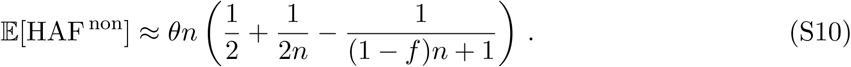

Because of the separation between carriers and non-carriers, the HAF-scores can be used to predict the carrier of ongoing selective sweeps without knowledge of the favored allele ^6^. Moreover, for the favored allele *e* with *fn* descendants, in a hard selective sweep that is not very close to fixation, we can approximate *ϕ*(*e*) as

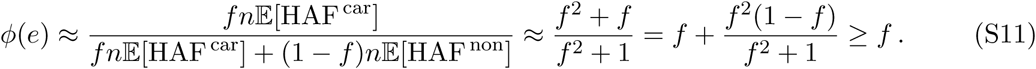

For a population undergoing a positive natural selection with favored mutation *e, ϕ*(*e*) overestimates the favored allele frequency *f* (Fig. 1B,E and Eq. S11). On the other hand, *κ*(*e*) underestimates *f* (Fig. 1B,E). Therefore, we expect the distribution of (*ϕ - κ*) for the favored allele to be skewed in positive direction.

SAFE score performs very well in separating the favored variant within a small window (See Fig. S4-S10); but the performance decays in larger windows (Fig. S8); because in larger windows most of the haplotypes become unique and *κ* estimate *f* correctly, even for favored mutations of selective sweeps, while we expect it to underestimate the *f* for the favored mutations. Consequently, the estimator *κ* is no-longer useful for pinpointing the favored mutation.

#### 1.4 Illustration of iSAFE: integrated SAFE for large regions

We devise iSAFE-score by extending the SAFE score to boost the performance in larger windows.We apply the SAFE score, as a kernel, on overlapping sliding windows. Define *S* as the set of all SNPs, *W* as the set of all sliding windows. Let *S*_1_ *v S* denote the subset of mutations that had the highest SAFE-score in their respective windows. For mutation *e ∈ S*, and window *w ∈ W*, let Ψ_*e,w*_ denote the SAFE-score of *e*, when *e* is ‘inserted’ into window *w* if it is positive, 0 otherwise. Fig. S3 provides a cartoon illustration of windows *w*_1_, *w*_2_, *w*_3_ and ⋆, ▴, and ▪, where ⋆ denotes the favored mutation and is located in *w*_2_.

We note the following:

- Ψ⋆,_*w*2_ is high for the favored mutation ⋆ However, Ψ▴, _*w*1_ and Ψ▪,_*w*3_ may be high even for hitchhiking mutations (▴,▪) due to the genealogies of *w*_1_ and *w*_3_. Thus SAFE-score by itself may not be a reliable predictor over a large region containing multiple windows.
- When a non-favored mutation is inserted in a window with a different genealogy, it is not likely to have a high SAFE-score. When ⋆and ▴ are inserted into window *w*_3_, Ψ⋆,_*w*3_> Ψ▴, _*w*3_ because ⋆ separates carriers from non-carriers and has high values for *ϕ*(⋆) and low values for *κ*(⋆). On the other hand, *κ*(▴) is higher because its descendants include non-carriers which are typically distinct haplotypes. Similarly Ψ⋆,_*w*1_ > Ψ▪,_*w*1_ because *ϕ* (▪) is lower in *w*_1_. In other words, the weighted sum of Ψ⋆,_*w*_ over all windows *w* is likely to dominate other mutations.
- Similarly, the window containing the favored mutation (*w*_2_) has the appropriate genealogy, and is likely to give a high score to multiple candidate mutations.

**Figure S3:**
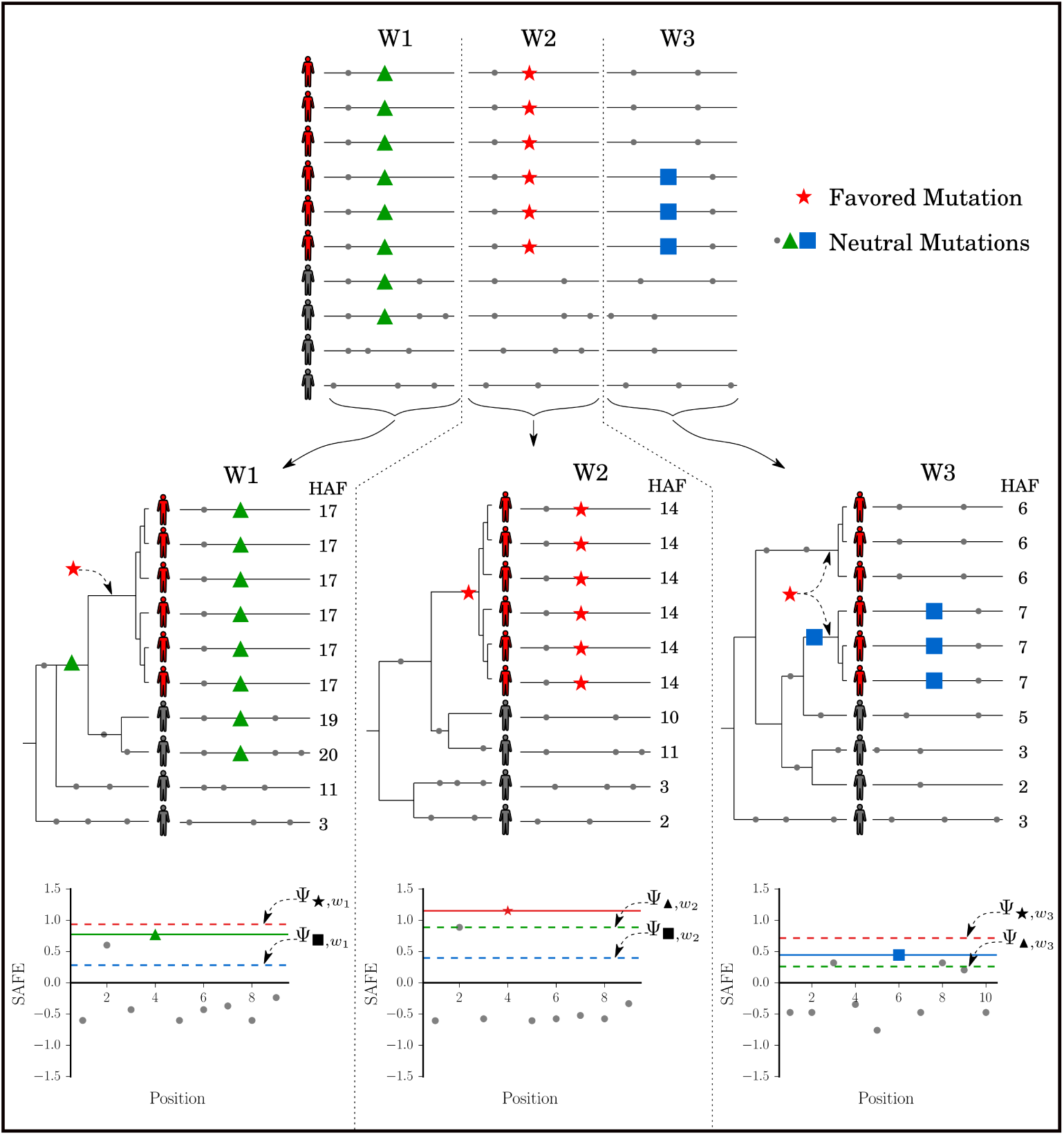
Cartoon illustration of iSAFE scoring. Window *w*_2_ carries the favored mutation (red ⋆) and also has a genealogy that separates carriers from non-carriers. Carrier haplotypes have a higher HAF-score. Windows *w*_1_, *w*_3_ do not have a mutation in their genealogies to separate carriers from non-carriers.

Based on these considerations, we define the score *α* of window *w ∈ W* as:

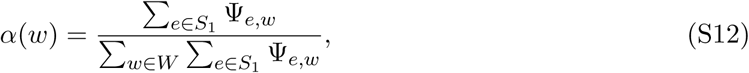

The window with the highest weight is the one which gets higher SAFE-scores for other mutations that are insrted into it. Finally, we define the score iSAFE of mutation *e ∈ S* as:

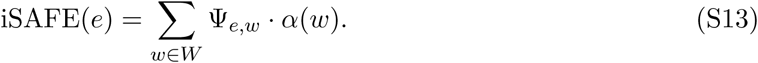

where the mutation with the highest score is one that gives high scores when inserted into high weight windows.

#### 1.5 MDDAF: <u>M</u>aximum <u>D</u>ifference in <u>D</u>erived <u>A</u>llele <u>F</u>requency

We have shown that iSAFE is successful in pinpointing the favored variant in an ongoing selective sweep. When the favored mutation is near fixation (*ν* > 0.9), iSAFE performance decays and when the favored variant is fixed (*ν* = 1), iSAFE cannot detect the favored mutation because it is no longer a variant (Fig. 3A). For the purpose of pinpointing the favored mutation in a fixed selective sweeps we add random samples from non-target population (outgroup) to the target population to constitute 10% of the sample.

To minimize the noise added to the data with random outgroup samples, we devise a simple method to decide whether to use outgroups or not. Our score is motivated by the work of Grossman et al.(2010) ^4^, who introduced the ΔDAF score of a mutation as 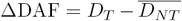, where *D*_*T*_ is the derived allele frequency in the target population and %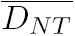 is the *average* derived allele frequency in non-target populations. As it is possible that some of the non-target populations are also under selection, choosing the average derived allele frequency may lower ΔDAF, and weaken the signal of selection. Instead we define the <u>M</u>aximum <u>D</u>ifference in <u>D</u>erived <u>A</u>llele <u>F</u>requency (MDDAF) score as:

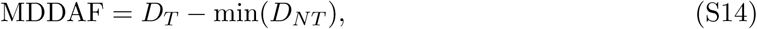

where, *D*_*T*_ is the derived allele frequency in the target population and min(*D*_*NT*_) is the *minimum* derived allele frequency over all non-target populations.

#### 1.6 Adding Outgroup Samples

Simulation of human population demography under neutral evolution (Fig. S20), shows *P* (MDDAF > 0.78*|D*_*T*_ > 0.9) = 0.001 (Fig. S22) making it a rare event to have high MDDAF score even when the frequency is high in the Target population. Therefore, when there is a high frequency mutation (*D*_*T*_ > 0.9) with MDDAF > 0.78 in the target population, we add random outgroup samples to the data to constitute 10% of the data. For analysis on real data, where we looked at 1000GP populations, we randomly selected outgroup samples from non-target populations of 1000GP.

In Fig. S16, we compared the performance of iSAFE with or without having the option of using outgroup samples; we simulated 5Mbp of human genome based on the human demography model described in Fig. S20. The selection happens in a random time, with a distribution given in Fig. S21, after the out of Africa in the lineage of EUR population (as the target population). When the onset of selection is before split of EUR and EAS (> 23kya), both (EUR and EAS) are under selection. When we have random sample option, we use the MDDAF criterion to decide whether we should use random sample or not. In case of adding random sample, we add a random subset of individuals from EAS+AFR to constitute 10% of the data (200 haplotypes from EUR and 22 from EAS+AFR).

The performance of iSAFE for sweeps with *ν* < 0.9 did not change with or without having outgroup sample option (Fig. 3A). When frequency of the favored mutation is near fixation (*ν* > 0.9) having the outgroup sample option is helpful and increase the performance of the iSAFE. When the sweep is fixed (*ν* = 1), iSAFE is no longer capable of detecting the favored mutation without having outgroup samples because the favored mutation is no longer a variant in the target population. However, with the outgroup sample option, iSAFE can successfully pinpoint the Favored mutation even in a fixed selective sweep (see Fig. 3A).

### 2 Simulation Experiments

#### 2.1 Default simulation parameters

Neutral and sweep samples were generated using the simulator *msms* ^14^. By default, simulated populations are haploid with sample size of *n* = 200 haplotypes from a larger effective population of *N* = 20000 haplotypes, each of length *L*, with default value 50kbp for SAFE and 5Mbp for iSAFE. For human populations, a mutation rate of approximately *µ* = 2.5 *·* 10-^8^ mutations per bp per generation^15,16^, and a recombination rate of approximately *r* = 1.25 *·* 10-^8^ per bp per generation ^17^ have been proposed. For SAFE simulations, we used a scaled mutation rate *θ* = 2*µN* = 1 mutations per kbp per generation and scaled recombination rate *ρ* = 2*rN* = 0.5 crossovers per kbp per meosis to approximate human rates. The rates were scaled linearly by *L*. In the case of positive selection the default scaled selection strength of the favored allele was set to *Ns* = 500, with the favored mutation located at a random position uniformly distributed on the range [1, *L*]. The default value for favored mutation starting frequency *v*_0_ = 1/*N* (hard sweep), and the frequency of the favored mutation (*ν*) at the time of sampling is a random value uniformly distributed on the range [0.1, 0.9]. We used the default parameters for all simulations unless otherwise stated.

#### 2.2 A model of human demography

We simulated demography of AFR, EUR, EAS populations with parameter shown in the Fig. S20 based on a popular demographic model of human population^18^. In case of positive selection, selection coefficient was set to *s* = 0.05 and starting favored allele frequency *v*_0_ = 0.001. The time of onset of selection was chosen at random (using the distribution in Fig. S21) after the out of Africa event, in the lineage of EUR population (as the target population). When the onset of selection is before split of EUR and EAS (> 23kya), both (EUR and EAS) are under selection.

### 3 Human Population Datasets

We downloaded the phased haplotypes of the 1000 Genomes Project (Phase 3; GRCh37) dataset from http://ftp.1000genomes.ebi.ac.uk/vol1/ftp/release/20130502/. The Ancestral Alleles dataset (GRCh37) is downloaded from http://ftp.ensembl.org/pub/release-75/fasta/ancestral-alleles/. The physical position was converted into genetic position using the genetic map in ftp://ftp-trace.ncbi.nih.gov/1000genomes/ftp/technical/working/20110106-recombination-hotspots/.

### 4 Computing Selection Statistics

#### 4.1 Computing iHS scores

We used the *selscan* ^19^ (v1.1.0a) software available at https://github.com/szpiech/selscan, with default settings to calculate the raw iHS ^8^ score. Next, we normalized the iHS score by estimating the distribution of raw iHS scores on 1,000 neutral simulations with the same simulation parameters. The iHS scores were always computed on a 5Mb window. When comparing results with SAFE on a 50kbp window, we used the corresponding iHS scores in the identical 50kbp region surrounding the favored variant (Fig. 1,S4). In considering 5Mb windows (Fig. S13), we compared the iHS scores on all variants for iHS against iSAFE.

#### 4.2 Computing SCCT scores

We used the SCCT (v1.1) software available at https://github.com/wavefancy/scct, provided by Wang et al. ^7^, with flanking SNPs size 300, and frequency interval 0.01.

#### 4.3 Computing CMS scores

CMS requires a control population as well as a demographic model in addition to the target population under selection. All CMS comparisons on simulated data were performed using a model of human demography^18^, described in Fig. S20, with a random onset of selection (Fig. S21). We used the CMS (v2.0) software available at https://github.com/broadinstitute/cms, disabling CMS’ default allele frequency filter in order to allow a more direct comparison with iSAFE SNP rankings.

### 5 Empirical *p*-val computation

We applied iSAFE on a neutrally evolving simulated population with window size 5Mbp, based on European demography shown in Fig. S20. A *p*-value was calculated based on empirical distribution of iSAFE on these simulated populations. We limited the number of samples to *∼*74,800,000 for efficiency, and this allows us to get a *p*-value as low as 1.34e-8 for iSAFE-score 0.304. Scores higher than this cut-off are considered to have *p*-value < 1.34e-8.

**Figure S4:**
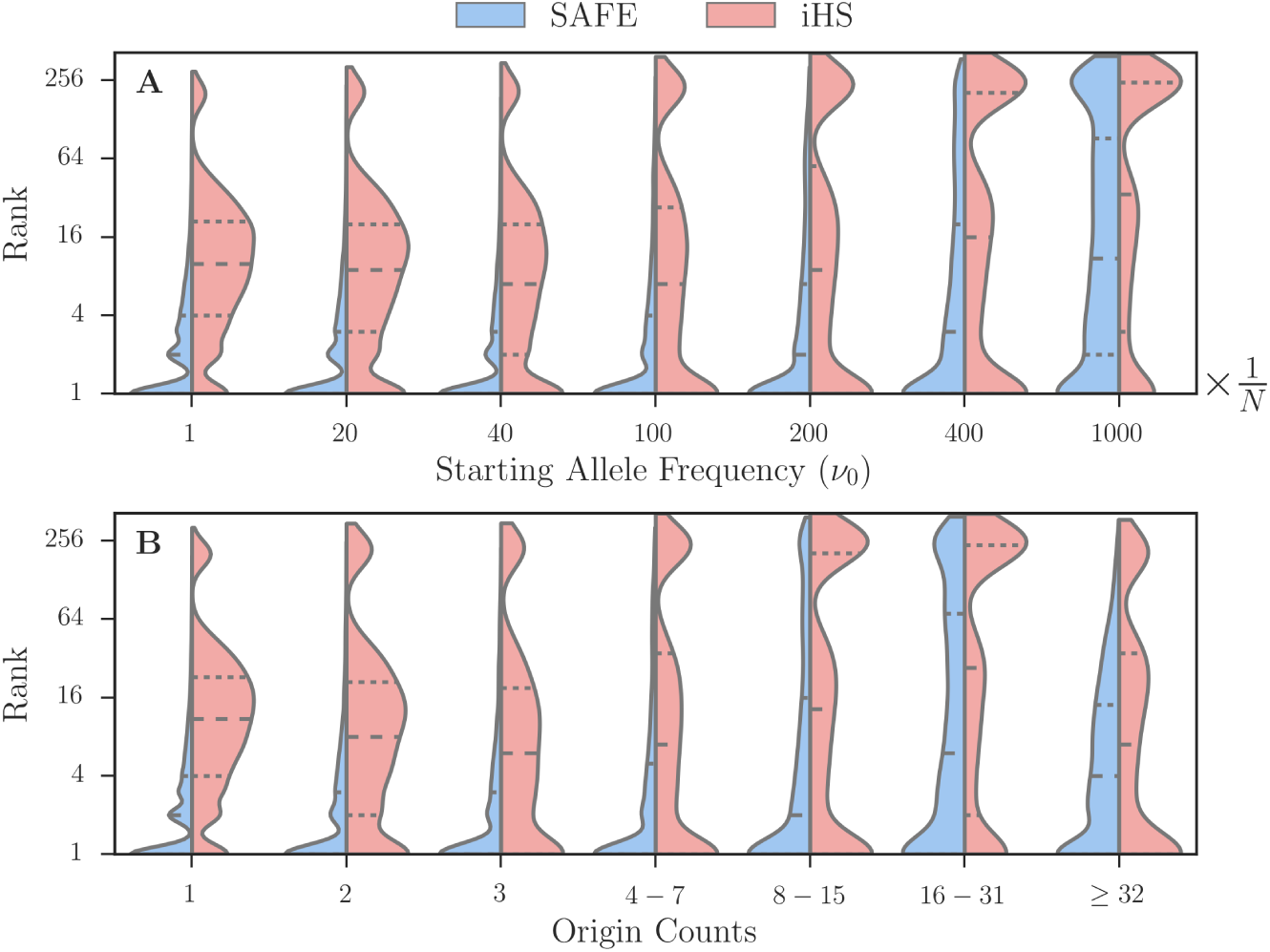
Performance of SAFE score on hard and soft sweeps. (**A**) Rank of the favored mutation for hard sweep (*v*_0_ = 1/*N*) and soft sweep (*v*_0_ > 1/*N*) in 1000 simulations per bin on 50kbp window with selection strength (*Ns* = 500) and fixed population size (*N* = 20, 000) and default values for other simulation parameters. The line with large dashes represents the median rank. (**B**) Rank of the favored mutation as a function of *Origin Count* (number of ancestors of carriers of favored Allele at the onset of selection pressure) for the same data as in A. Origin Count of hard sweep is always 1.

**Figure S5:**
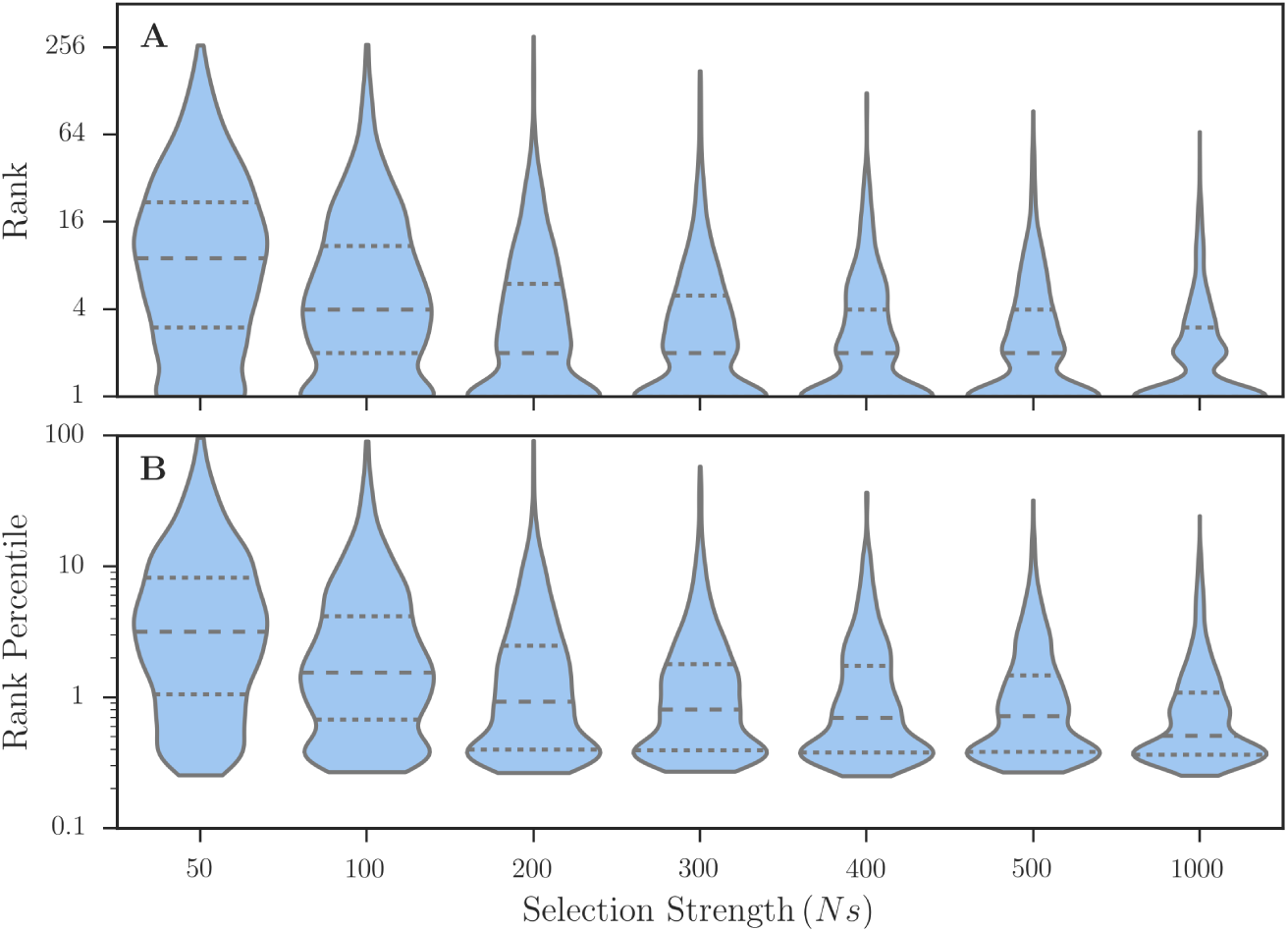
Performance of SAFE score with different selection strength. (**A, B**) Rank and rank percentile of the favored mutation as a function of selection strength (*Ns*) in 1000 simulations per bin on 50kbp window with default values for other simulation parameters. The line with large dashes represents the median rank.

**Figure S6:**
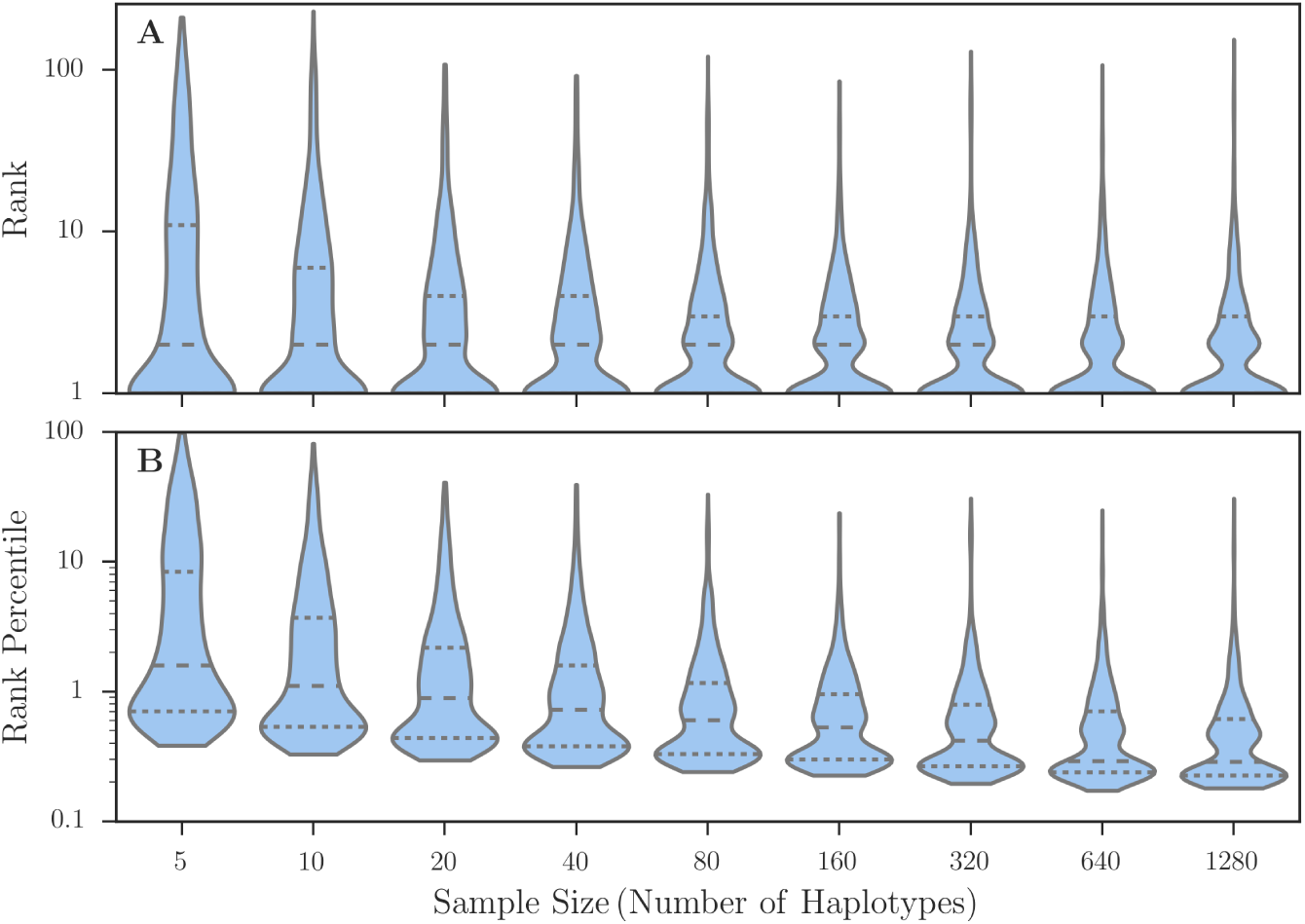
Performance of SAFE score with different sample size. (**A, B**) Rank and rank percentile of the favored mutation as a function of sample size in 1000 simulations per bin with selection strength (*Ns* = 500) and default values for other simulation parameters. The line with large dashes represents the median rank.

**Figure S7:**
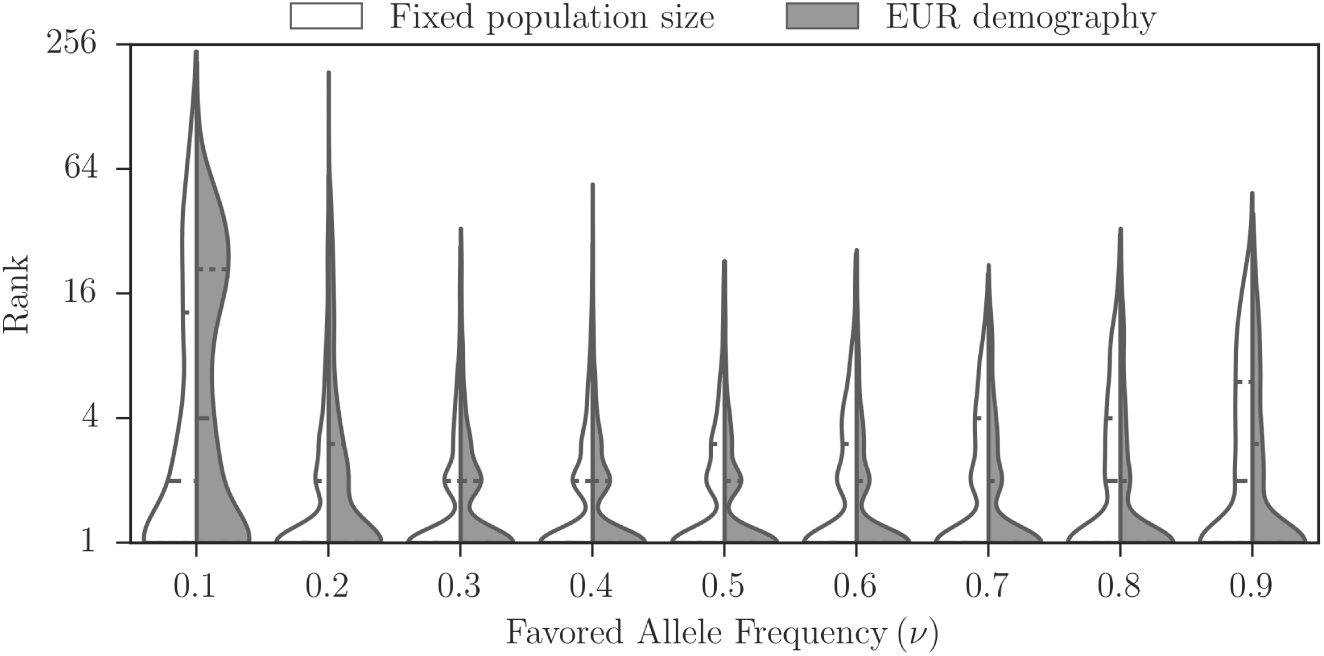
Performance of SAFE score in a model of European demography. White represents the result for a fixed size population model with default parameters and gray represents a model of human demography for EUR population. The model and all the parameters used are described in Fig. S20. The onset times of selection was post-bottleneck (23 kya-current) epochs. 1000 samples per bin were simulated with default values for simulation parameters not assigned in Fig. S20.

**Figure S8:**
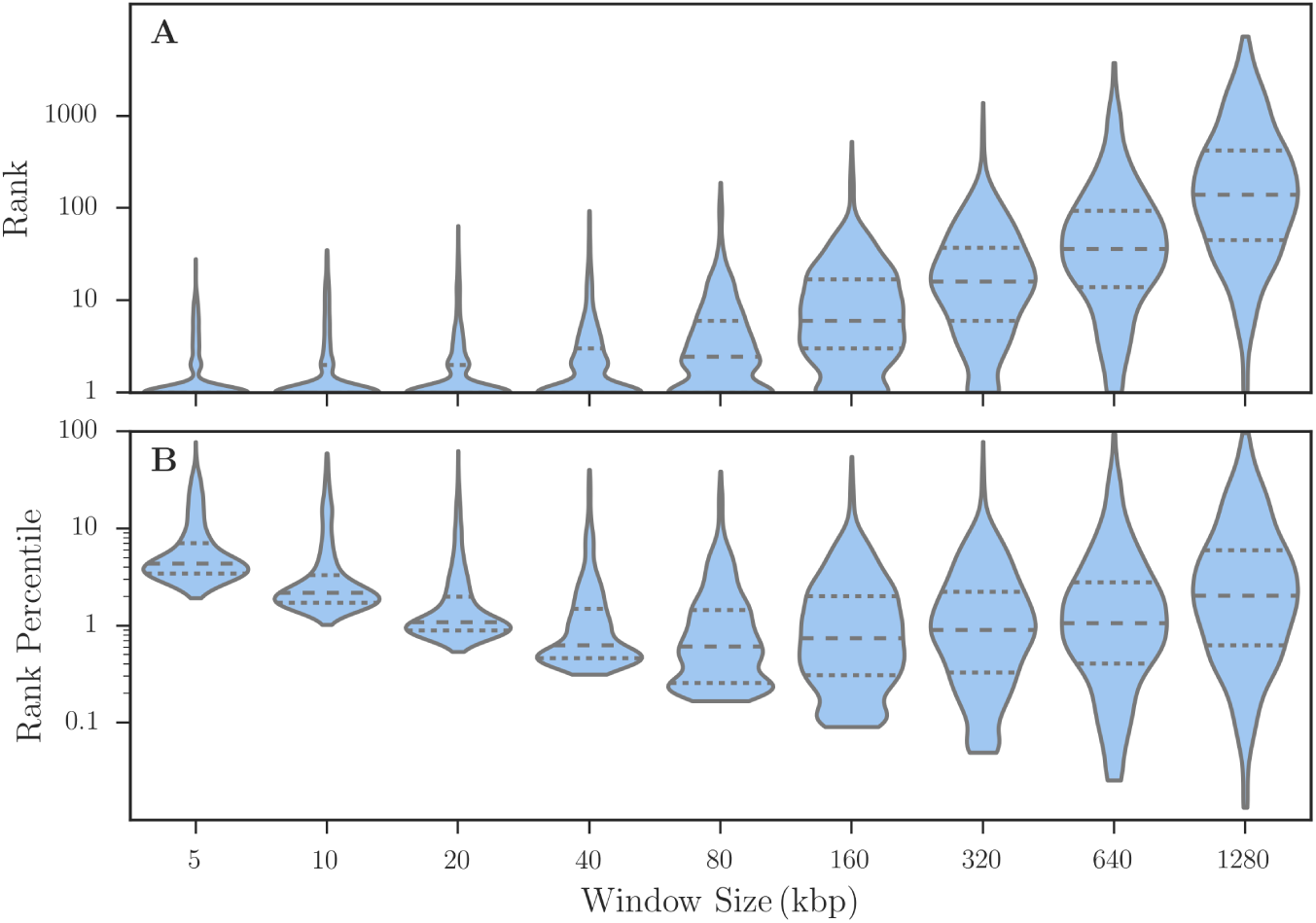
SAFE & window size. (**A, B**) Rank and rank percentile of the favored mutation as a function of window size in 1000 simulations per bin with selection strength (*Ns* = 500) and default values for other simulation parameters. The line with large dashes represents the median rank.

**Figure S9:**
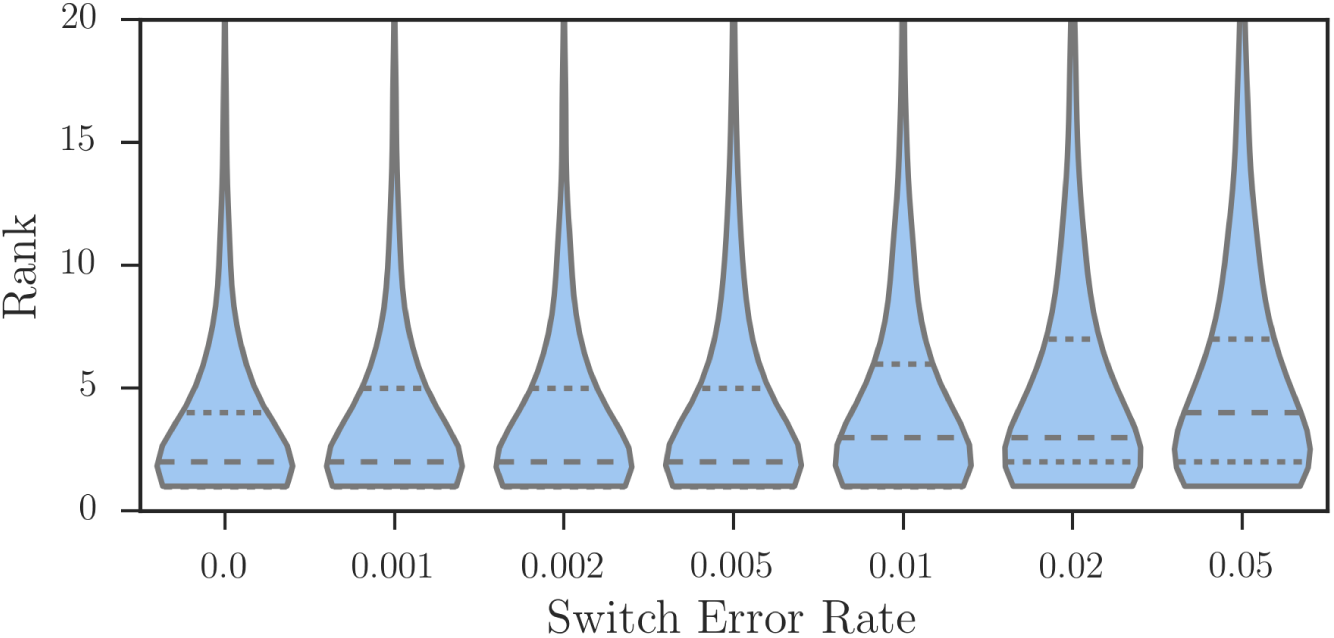
Performance of SAFE score with different switch error rate. Rank of the favored mutation as a function of haplotype phasing switch error rate in 1000 simulations with selection strength (*Ns* = 500) and default values for other simulation parameters. The line with large dashes represents the median rank.

**Figure S10:**
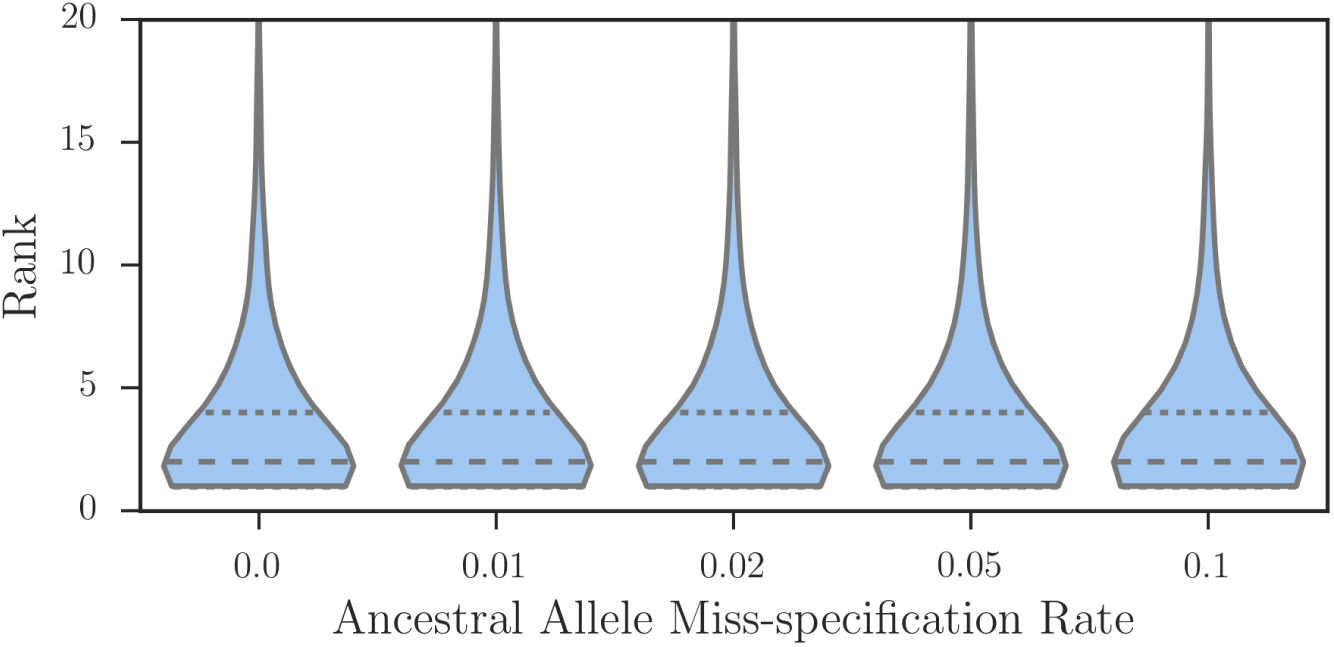
Performance of SAFE score with different rate of ancestral allele miss-specification. Rank of the favored mutation as a function of ancestral allele miss-specification rate, given that the ancestral allele of the favored mutation is specified correctly, in 1000 simulations with selection strength (*Ns* = 500) and default values for other simulation parameters. The line with large dashes represents the median rank.

**Figure S11:**
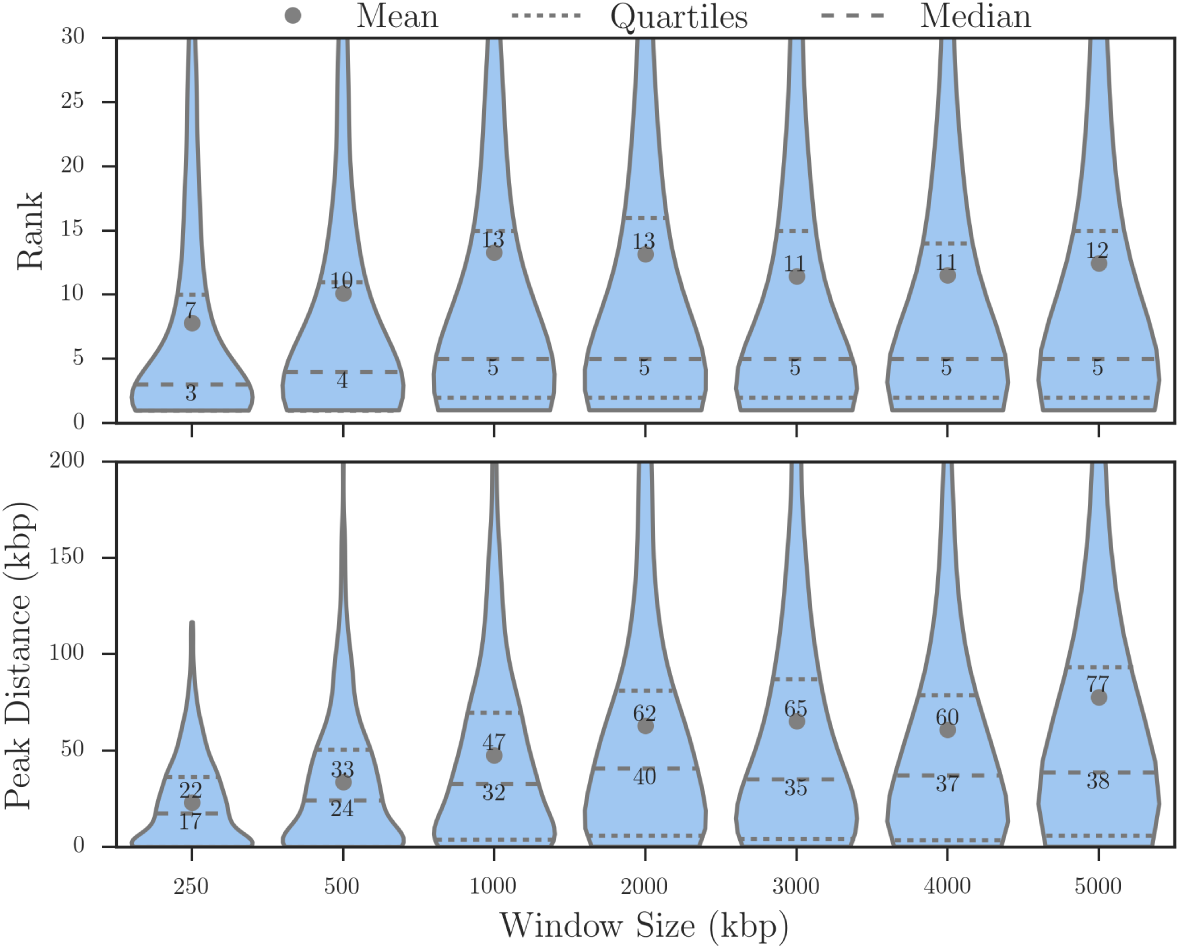
iSAFE and Window Size. Performance of iSAFE measured by rank of the favored variant and the distance of the favored variant from the peak in 1000 simulations per bin. The line with large dashes represents the median rank.

**Figure S12:**
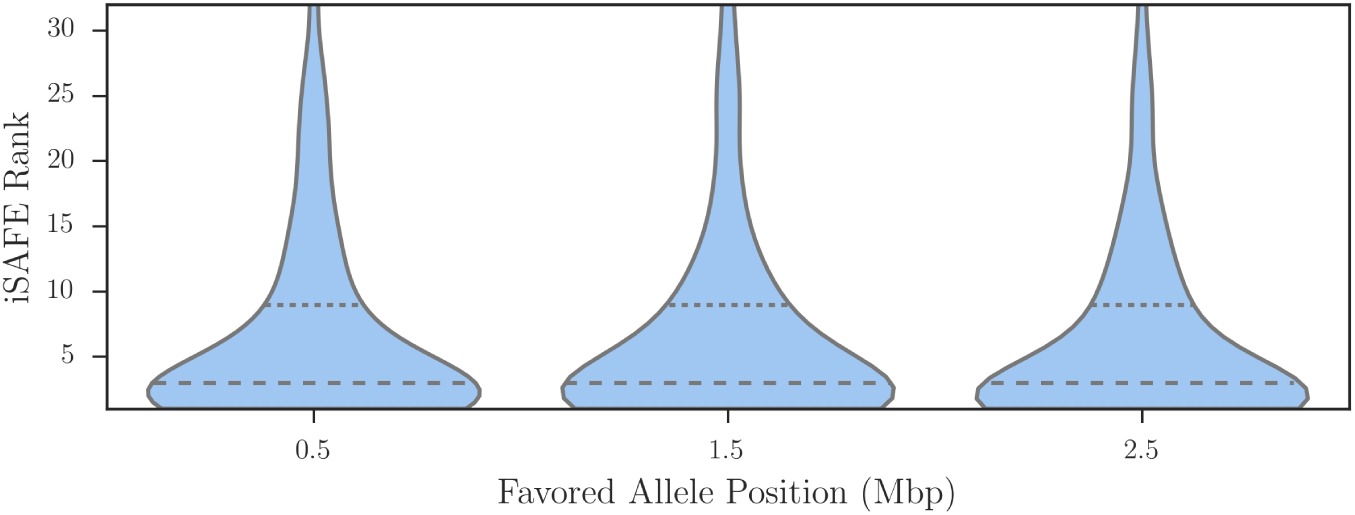
iSAFE and Position of the Favored Mutation. iSAFE rank of the favored mutation on 5Mbp regions with different position of the favored mutation. Each bin includes 1000 simulations with the position of the favored mutation selected from [0.5Mbp, 1.5Mbp, 2.5Mbp]. The dashed (dotted) line represents median (upper quartile). This result shows iSAFE is robust to the position of the favored mutation.

**Figure S13:**
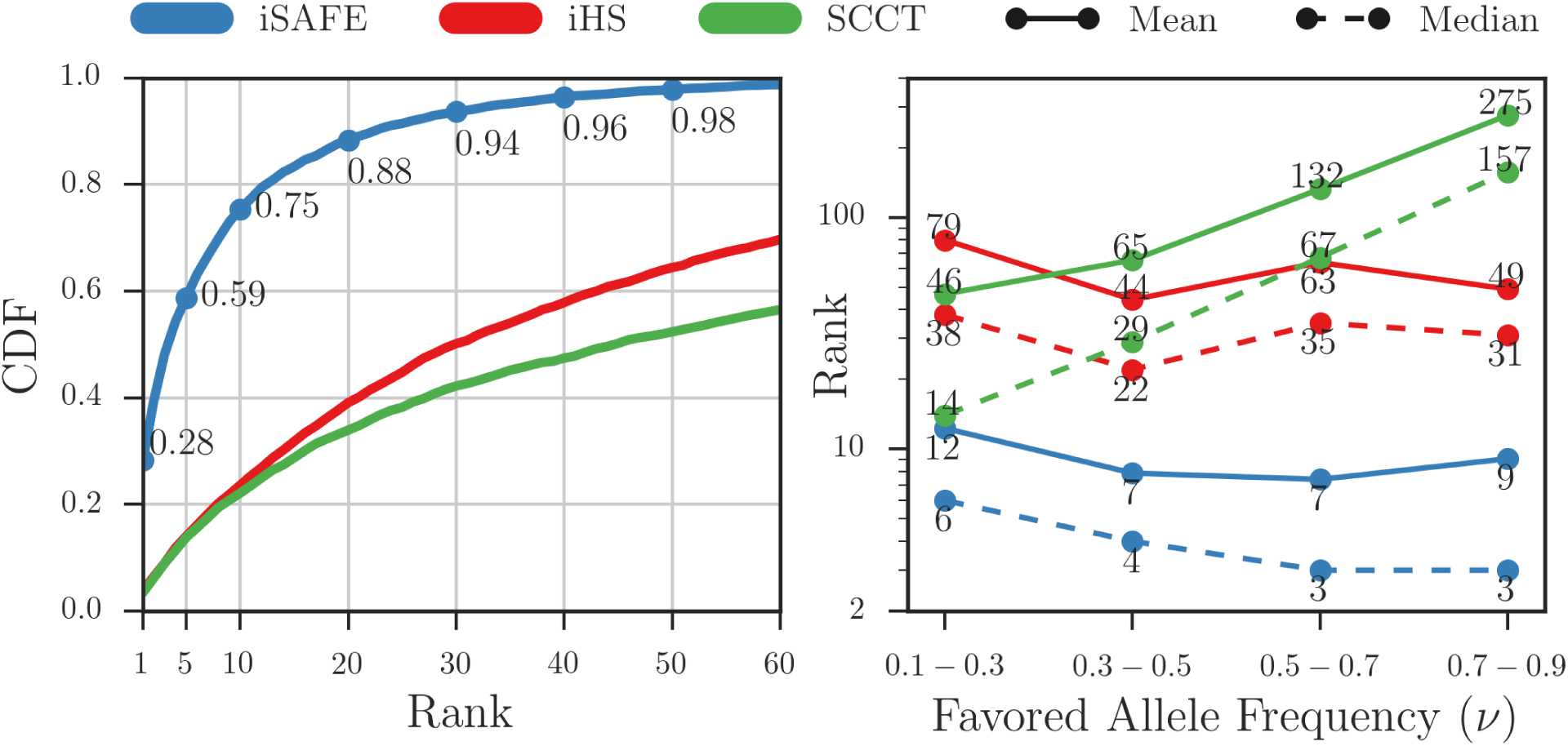
iSAFE compared to iHS and SCCT. Performance of iSAFE compared to iHS and SCCT measured by rank of the favored variant in 5000 simulations on 5Mbp region around ongoing hard sweeps (*v*_0_ = 1/*N*, 0.1 < *v* < 0.9) with a fixed population size (*N* = 20, 000) and default values for other simulation parameters. In the left panel, for any rank *r* on the X-axis, the *y*-intercept represents the proportion of samples where the favored allele had rank *≤ r*. In the right panel, solid (dashed) lines represent the mean (respectively, median) value of the favored allele rank.

**Figure S14:**
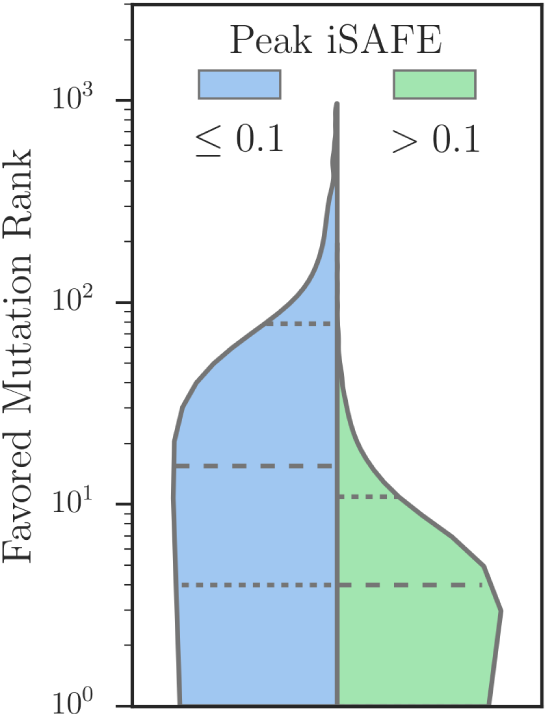
Peak iSAFE. Empirical analysis, with 5000 simulations on 5Mbp region with a wide range of selection strength (*Ns ∈* [10, 50, 100, 200, 300, 400, 500, 1000]), shows difference in performance of iSAFE beyond a score threshold of 0.1 for peak value of iSAFE (see Fig. 3C).

**Figure S15:**
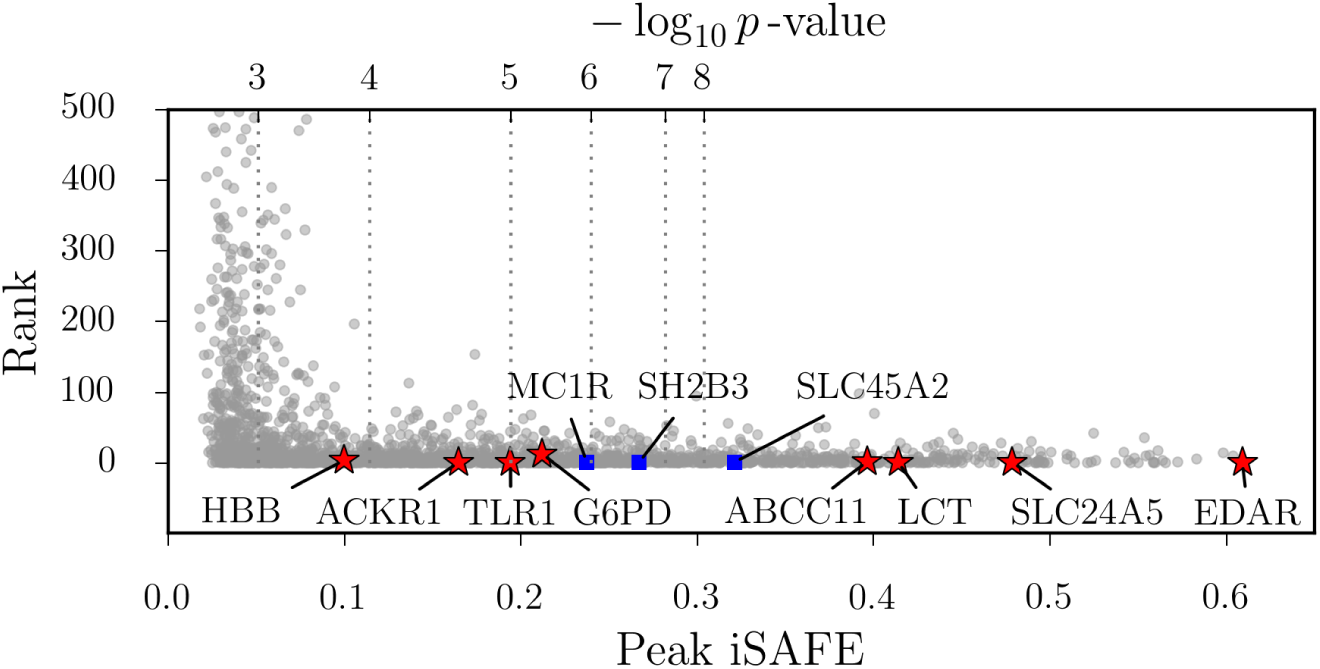
Peak iSAFE. Rank of favored mutation as a function of iSAFE-score (Bottom x-axis) or p-value (top x-axis). Each gray dot represents the favored mutation of a simulation using a wide range of selection coefficients. The performance deteriorates for iSAFE-scores below 0.1.

**Figure S16:**
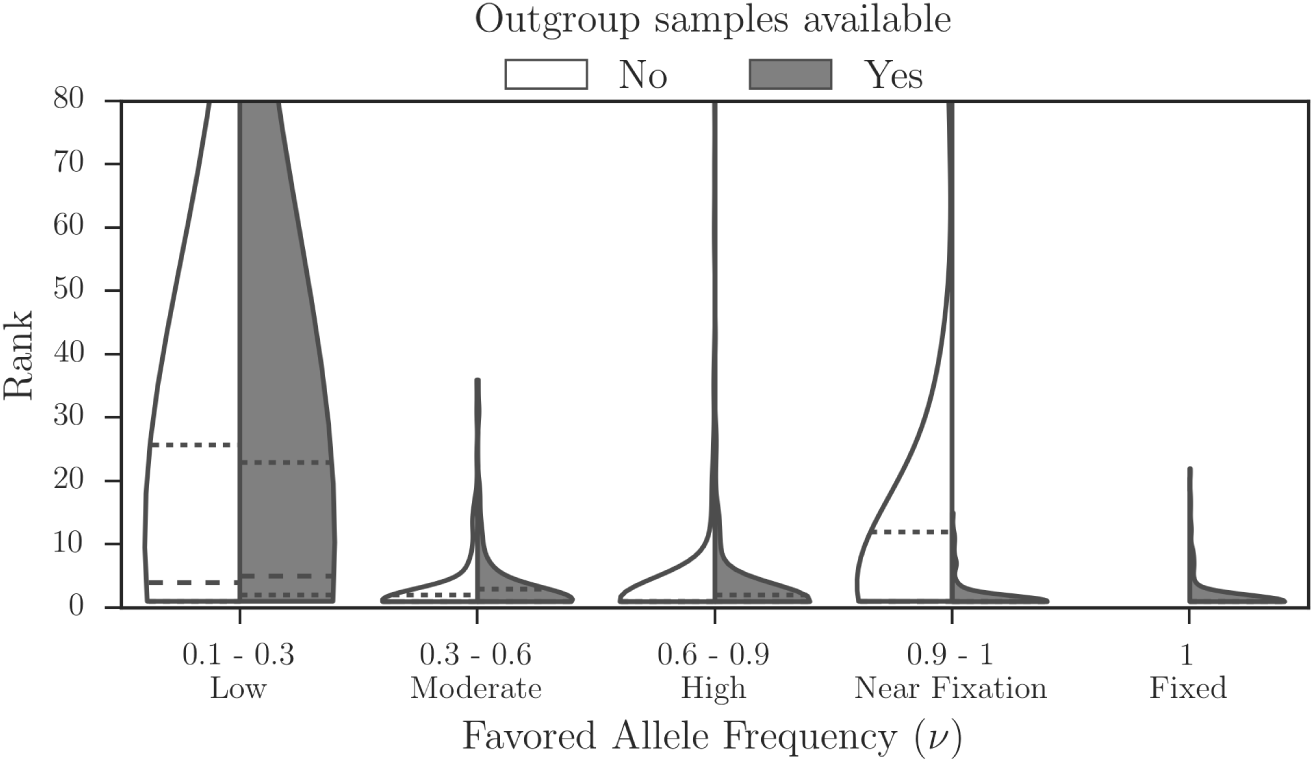
iSAFE and Outgroup Samples. iSAFE performance upon addition of outgroup samples. No deterioration is seen for low frequencies of the favored variant, but iSAFE performance improves dramatically when favored mutation is near fixation or fixed..

**Figure S17:**
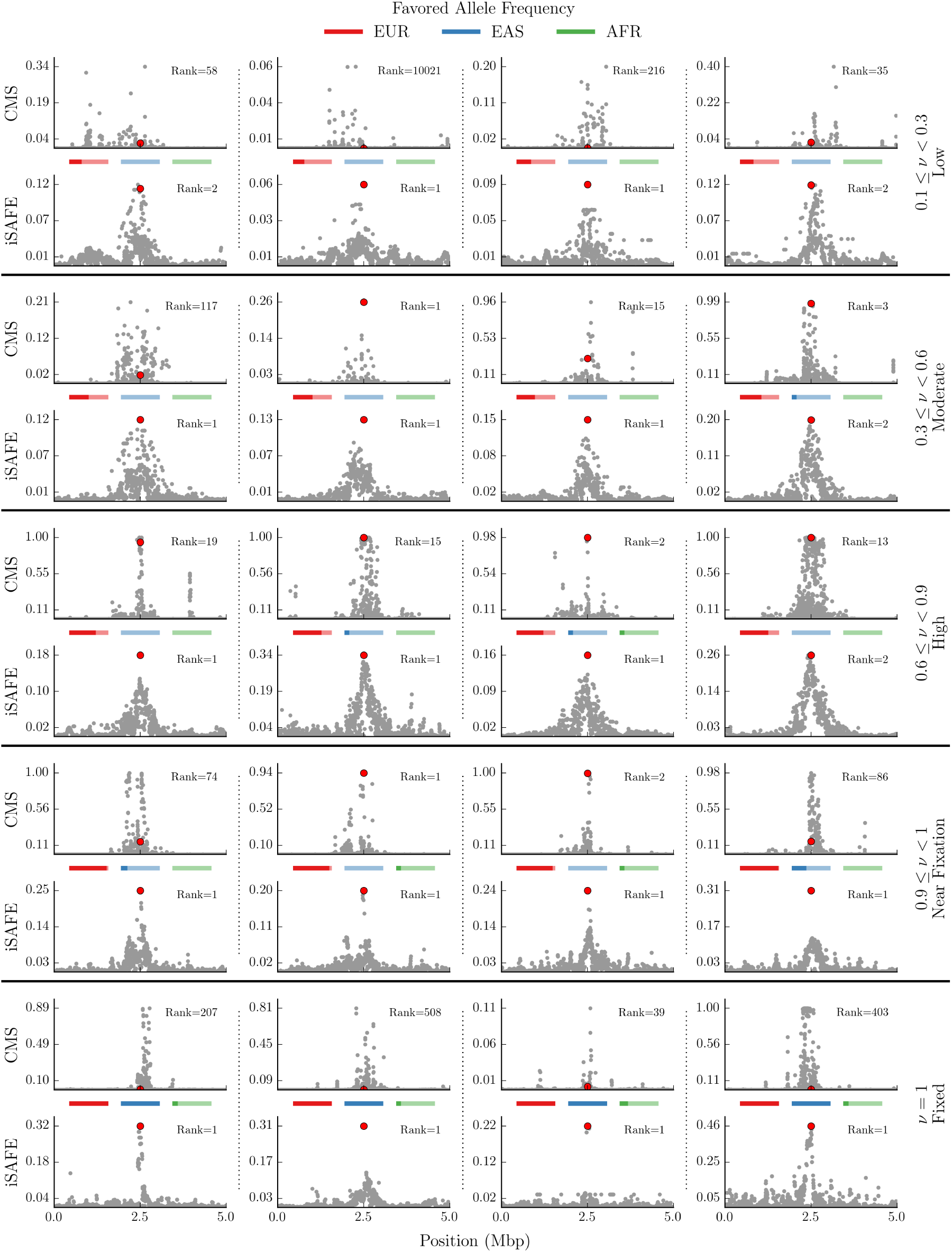
Demo I: iSAFE vs. CMS in a model of human demography. Comparing iSAFE and CMS signals in a model of human demography described in Fig. S20, S21. Solid-horizontal lines separate replicates based on the favored allele frequency (*v*) in EUR as the target population, and dotted-vertical lines separate different replicates. The rank of the favored mutation (solid-red circle) for each test is shown on the top-right corner.

**Figure S18:**
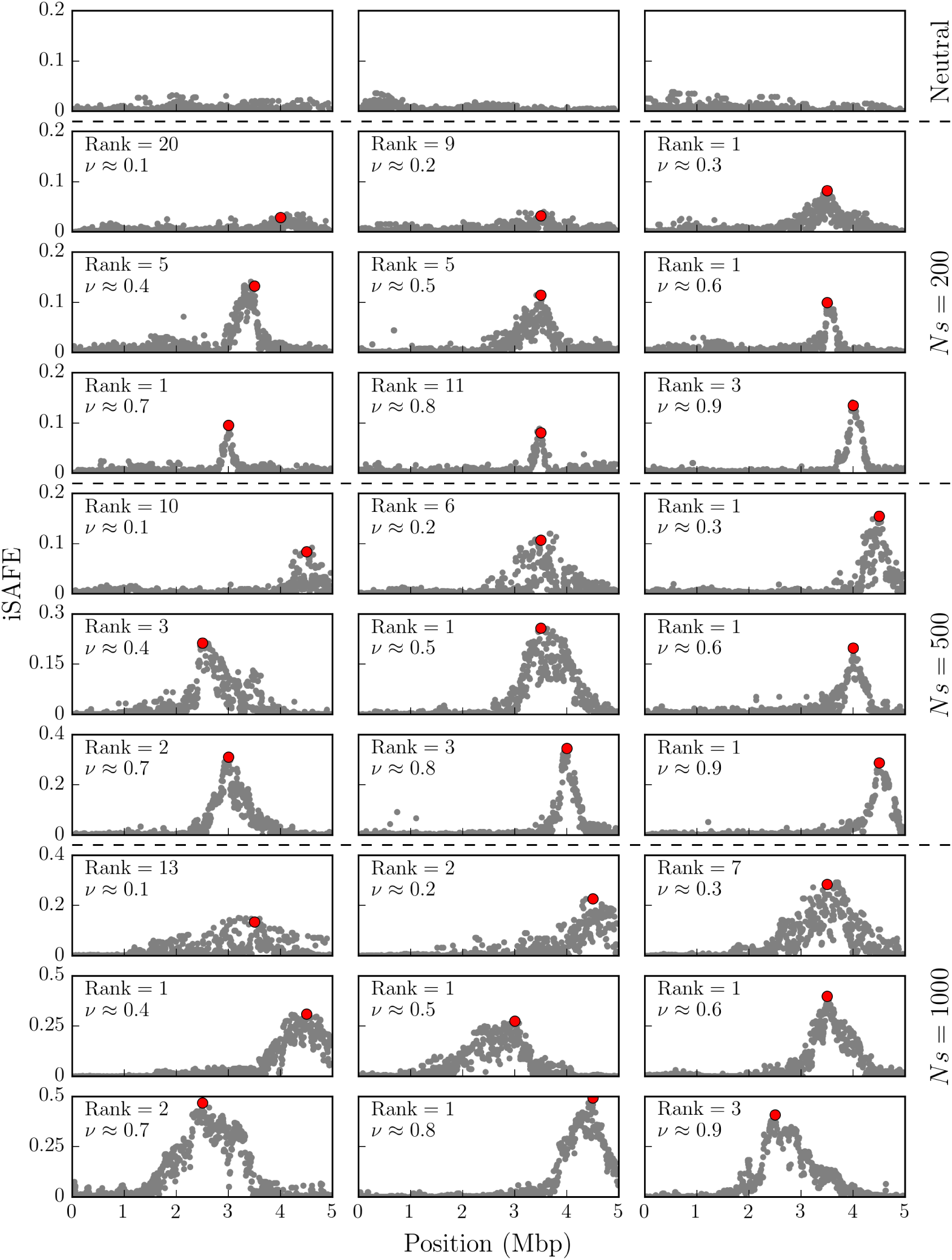
Demo II: iSAFE without outgroup samples. iSAFE on ongoing selective sweeps with different favored allele frequency (*v*) in 5Mbp region. The position of the favored mutation selected from range [2.5Mbp, 5Mbp]. Other simulation parameters are the default values for fixed population size and outgroup samples are not available.

**Figure S19:**
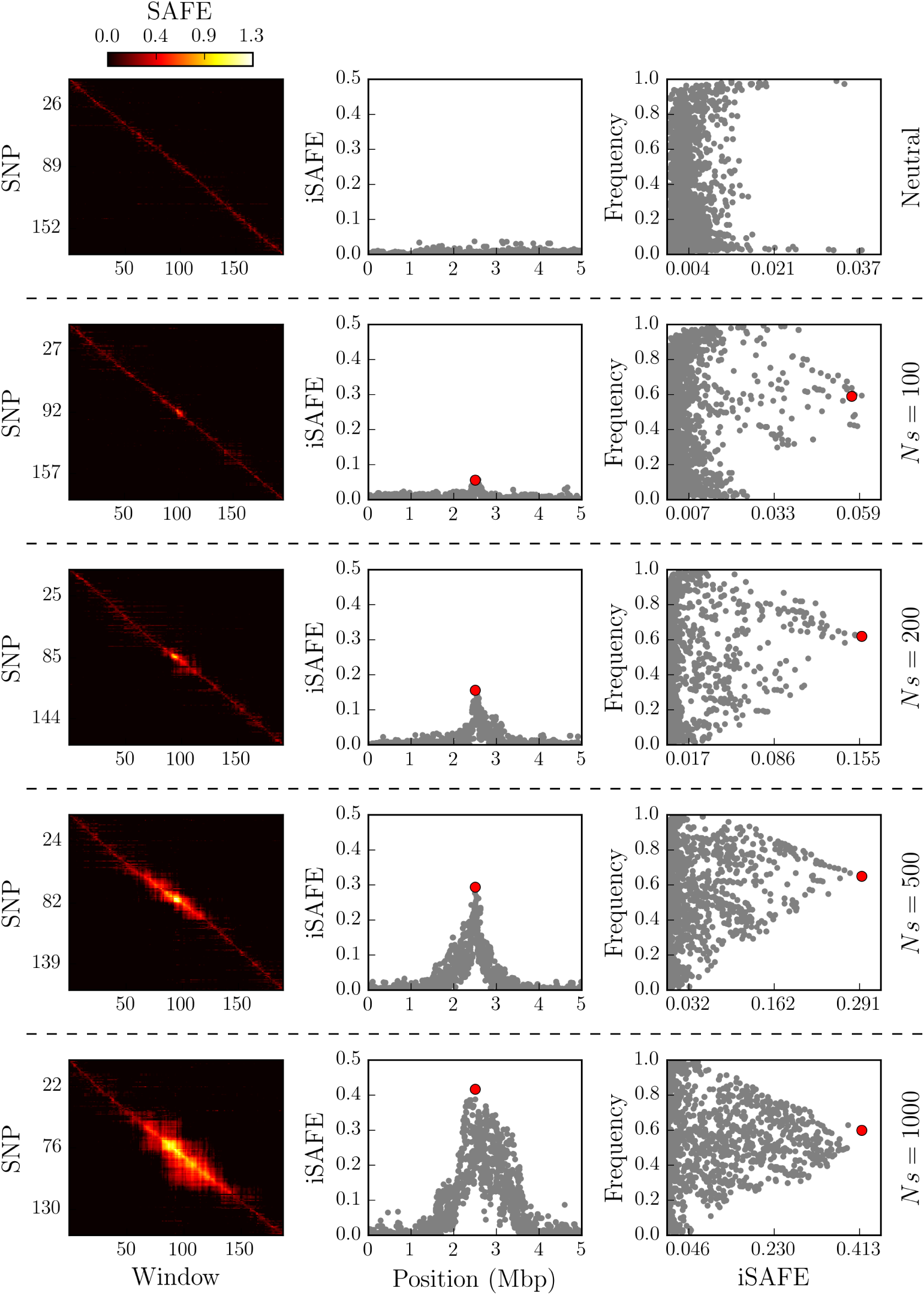
Demo III: iSAFE and selection strength. iSAFE on 5Mbp region with different selection strength, *Ns ∈* [0, 100, 200, 500, 1000]. Left panels shows the Ψ^1^ matrix. Middle panel shows the iSAFE-score as a function of the variant position. Right panel show the derived allele frequency as a function of iSAFE score.

**Figure S20:**
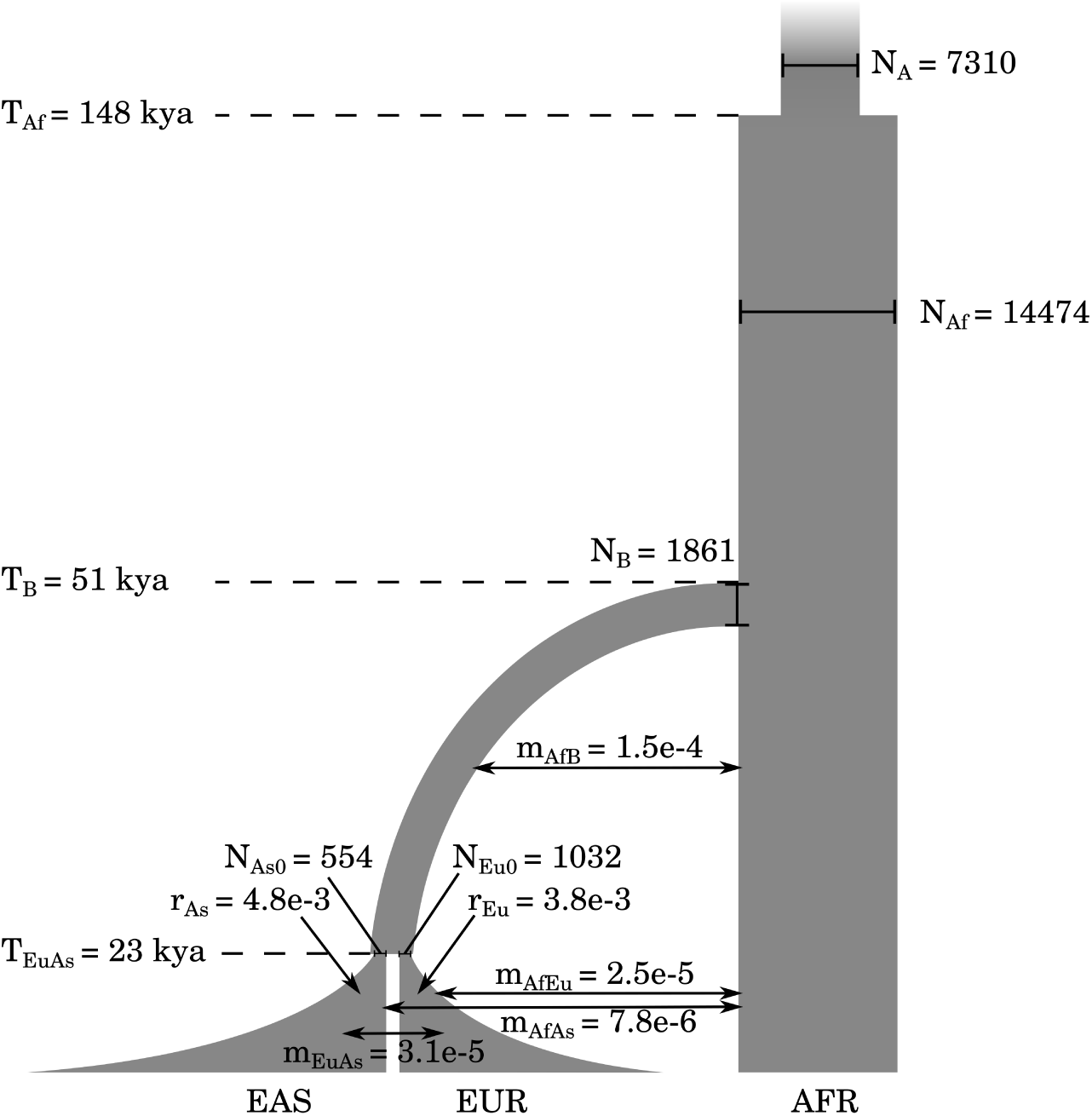
A model of human demography described by Fig. 4 and Table 2 of Gravel et al. (2011)^18^. The model assumes an out-of-Africa split at time *T*_*B*_, with a bottleneck that reduced the effective population from *N*_Af_ to *N*_*B*_, allowing for migrations at rate *M*_Af-B_. The African population stays constant at *N*_Af_ up to the present generation. The model assumes a second split between European and Asian populations at time *T*_EuAs_, with a bottleneck reducing the Asian and European populations to *N*_As0_ and *N*_Eu0_ respectively. The bottleneck was followed by exponential growth at rates *r*_As_ and *r*_Eu_, as well as migrations among all three sub-populations, leading to current populations from which East Asian (EAS), European (EUR), and Africans (AFR) individuals were sampled.

**Figure S21:**
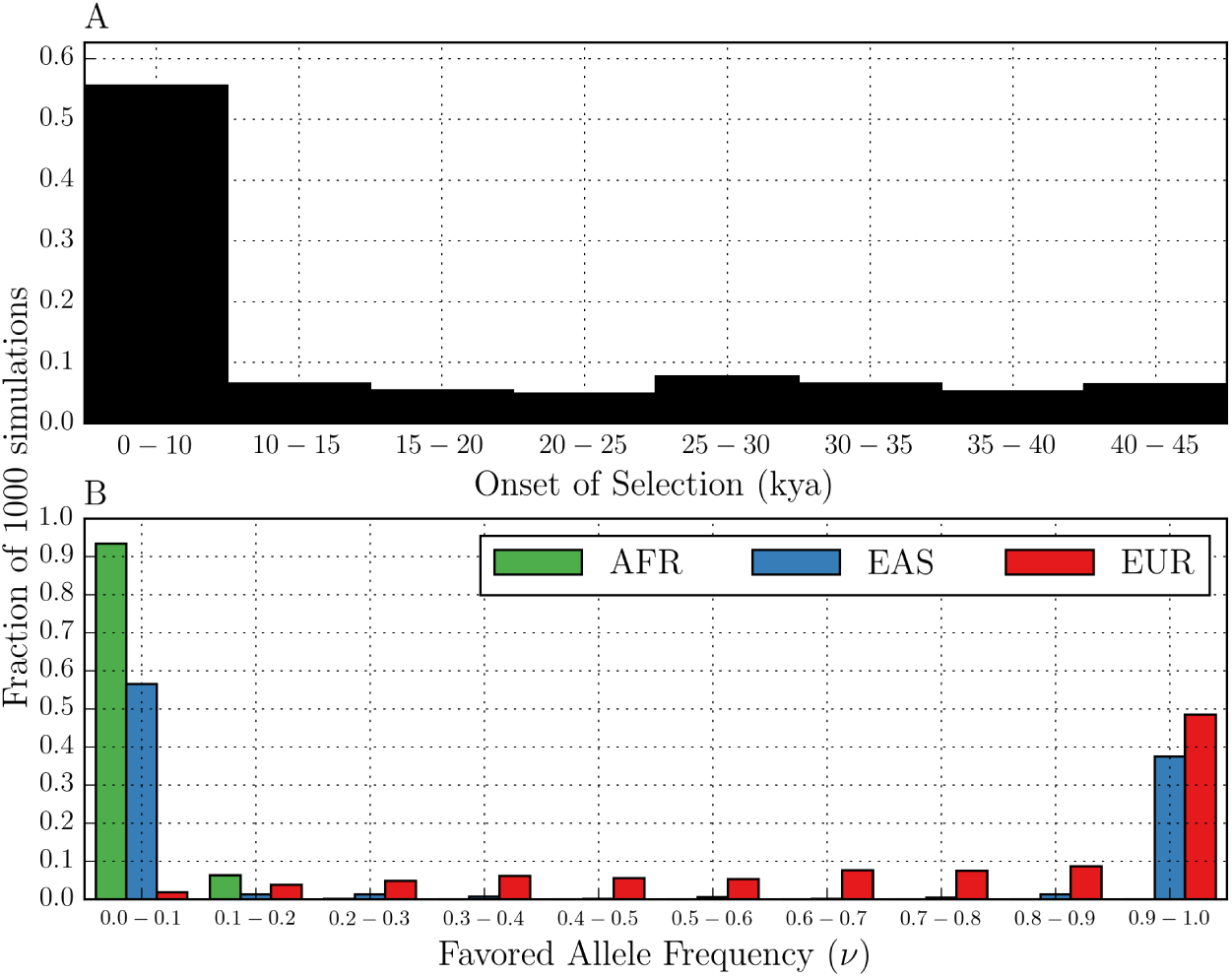
Simulation of Selection on Human Demography. We simulated 1000 selective sweeps on 5Mbp region based on the human demography model described in Fig. S20, and with selection coefficient *s* = 0.05 and starting favored allele frequency *v*_0_ = 0.001. **A)**The selection happens in a random time, after the out of Africa in the lineage of EUR population (as the target population). **B)**When the onset of selection is before split of EUR and EAS (> 23kya), both (EUR and EAS) are under selection.

**Figure S22:**
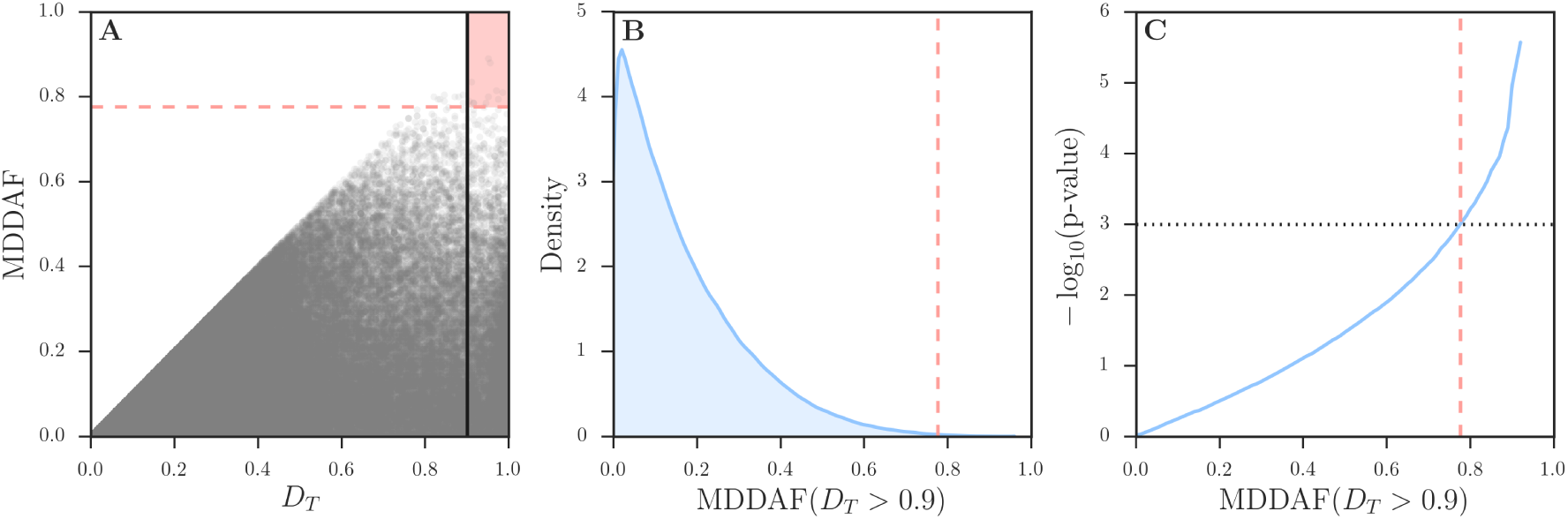
Maximum Difference in Derived Allele Frequency (MDDAF) . We simulated 25, 000 instances of AFR, EUR, and EAS populations, based on a demographic model described in Fig. S20. We used default values for simulation parameters not assigned in the Fig. S20. **A)**The MDDAF score of mutations as a function of derived allele frequency in the target population *D*_*T*_.**B)**Distribution of the MDDAF score for mutations with *D*_*T*_ > 0.9. **C)**P-value of the MDDAF score for mutations with *D*_*T*_ > 0.9. The dashed-red lines represent the value 0.78, where MDDAF, given *D*_*T*_ > 0.9, has a p-value less than 0.1%.

### 6 Results on selective sweeps in human populations

#### 6.1 Well characterized selective sweeps

We examined 8 well characterized selective sweeps with strong candidate mutation. These genes are LCT, SLC24A5, TLR1, EDAR, ACKR1/DARC, ABCC11, HBB, and G6PD^4,20,21,22,23,24^. iSAFE results for these genes are summarized in Fig. 3 and Table S1.

We also examined 14 other regions reported to be under selection with one or more candidate favored mutations^9,25,26,4,27^.

#### 6.2 Pigmentation genes

##### SLC45A2/MATP

This region is involved in human pigmentation pathways and is a target of selective sweep in European population^9^. A nonsynonymous mutation rs16891982 is associated with light skin pigmentation and is believed to be the favored variant^4,9^. This mutation is also ranked first by iSAFE out of *∼*21,000 mutations (5Mbp) in CEU population with a significant score (see Fig. 3N, iSAFE=0.32, *p*-val<1.3e-8).This mutation is almost fixed in European; frequency in AFR, EAS, SAS, AMR, and EUR is 0.04, 0.01, 0.06, 0.45, and 0.94, respectively.

##### MC1R

The MC1R gene is implicated in many skin color phenotypes, including red hair, fair skin, freckles, poor tanning response and higher risk of skin cancer. It is is a target of positive selection in East Asian populations, with a non-synonymous mutation (rs885479) suggested as a candidate favored mutation^25^. This mutation is ranked first by iSAFE in CHB+JPT (see Fig. 3P, iSAFE =0.24, *p*-val = 1.4e-6) out of *∼*16,000 mutations (2.8Mbp). The putative selected region is 300kbp away from the telomere of chromosome 16.

##### GRM5-TYR

The Tyrosinase (TYR) gene, encoding an enzyme involved in the first step of melanin production is present in a large region under selection. A nonsynonymous mutation rs1042602 in TYR gene is reported as a candidate favored variant^9^. A second intronic variant rs10831496 in GRM5 gene, 396kbp upstream of TYR, has been shown to have a strong association with skin color ^10^.

In contrast, iSAFE ranks mutation rs672144 as the top candidate for the favored variant region out of *∼*22,000 mutations (5Mbp). This variant was the top ranked mutation not only in CEU (iSAFE = 0.48, *p*-val*≪*1.3e-8), but also the top ranked mutation for EUR, EAS, AMR, and SAS (see Fig. 3Q and Fig. S23). The signal of selection is strong in all populations (iSAFE >0.5, *p*-val*≪*1.3e-8 for all of) except AFR, which does not show a signal of selection in this region. It may not have been reported earlier because it is near fixation in all populations of 1000GP except for AFR (*f* = 0.27), as seen in Fig. S23G. We plotted the haplotypes carrying rs672144 and found (Fig. 4) that two distinct haplotypes carry the mutation, both with high frequencies maintained across a large stretch of the region, suggestive of a soft sweep with standing variation.

The previously suggested candidates rs1042602, rs10831496 are fully linked to rs672144 (Fig. S24), but not to each other. The EUR haplotypes can be partitioned into 4 clusters (Fig. S24). Each of the 4 haplotypes show high homozygosity, suggestive of selection. However, rs1042602 can only explain the sweep in clusters C1+C2. rs10831496 can only explain C1+C3. Only rs672144 explains all 4 clusters, providing a simpler explanation of selection in this region. GTEx eQTL analysis on TYR gene for the tissue ‘Skin - Sun Exposed (Lower leg)’ showed *p*-value 0.61 for rs1042602, *p*-value 0.15 for rs10831496, and *p*-value = 0.08 for rs672144. While the *p*-value does not rise to a level of significance due to sample size issues, it is indicative of a regulatory function for the mutation.

##### OCA2-HERC2

This region is suggested as a target of selection in European^4,28,9^, and several mutations in this region are associated with hair, eye, and skin pigmentation. For example, rs12913832 is considered to be the main determinant of iris pigmentation (brown/blue) and is also associated with skin and hair pigmentation and the propensity to tan^9^. rs1667394 is also linked to blond hair and blue eyes ^28^. Some other mutations, many fully linked, (rs4778138, rs4778241, rs7495174, rs1129038, rs916977) are also associated with blue eyes^28^. This region is also suggested to be a target of selection in East Asia with rs1800414 suggested as a candidate for light skin pigmentation in that population. We applied iSAFE on this region to all 1000GP super-populations.

iSAFE selected a single variant rs1448484 in OCA2 (with high confidence, *p*-val<1.34e-8 for EUR, EAS, AMR and *p*-val=2.13e-6 for SAS) as the favored variant in all 1000GP populations (EUR, EAS, SAS, AMR) except for AFR that showed no signal of selection in this region (see Fig. S25 and Fig. 3P). This variant is close to fixation in all populations except for AFR, where *ν* = 20% (see Fig. S25F). iSAFE result along with the frequency pattern of the top ranked variant, suggests an out of Africa selection, probably on light skin color, on this region. The other candidate variants are all ranked high, and tightly linked with the top-ranked variant (Table S2).

##### KITLG

This genomic region has been linked to skin pigmentation^29^ in European and East Asian populations, and shows a strong signature of selective sweep on regulatory regions surrounding the gene in all non-African populations ^25^, with a candidate variant rs642742, that is associated with skin pigmentation^29^.

iSAFE analysis identified the same mutations gaining the top rank in multiple populations (Fig. S26). Top rank mutations in EUR, SAS, EAS, and AMR populations are shown in Table S3. The top ranked mutation in EUR and CEU populations (rs405647) was ranked 1, 2, 3 in AMR, SAS, and EAS, respectively, and is tightly linked to rs642742 (*D*^*t*^ = 0.92). Mutation rs661114 is ranked 2 in EUR, 5 in CEU, 6 in SAS, and 20 in AMR, and lies in a region with H3K27 acetylation that is associated with enhanced expression.

##### TRPV6

This region has been reported a target of selection in CEU population^26^. TRPV6 is involved in calcium absorption. It has been suggested that “Individuals with lighter skin pig-mentation might have produced too much 1,25-dihydroxyvitamin D, resulting in an increased intestinal Ca2+ absorption. Thus, to reduce the risk of absorptive hypercalciuria with kidney stones, the derived haplotype would have spread only among individuals with lighter skin pigmentation”^30^. iSAFE suggests 10 strongly linked mutations located along a 9kbp region located 84kbp downstream of TRPV6 (see Fig. S28). These mutations are ranked in the top 10 in all non-African populations (Table S5). There is no signal of selection in this region in AFR. The pattern of selection in this region in global population along with the confidence and consistency of iSAFE results in all non-African populations is consistent with an out of Africa selection on this region with the favored mutation being near fixation in all non-African populations (Fig. S27).

#### 6.3 Population specific selection: East Asian

##### PCDH15

This gene plays a role in development of inner-ear hair cells and maintaining retinal photoreceptors and is reported to be under selection in East Asian and a nonsynonymous mutation rs4935502 is proposed to be the favored variant^4^. This mutation is ranked 12 by iSAFE in CHB+JPT (see Fig. S30A, iSAFE =0.45, *p*-val<1.34e-8). All top mutations are highly linked.

##### ADH1B

”The ADH1B gene encodes one of three subunits of the Alcohol dehydrogenase (ADH1) protein, a major enzyme in the alcohol degradation pathway that catalyzes the oxidization of alcohols into aldehydes.” This region is a target of positive selection in East Asian population^26^. A non-synonymous mutation in this gene is associated with Alcohol dependence ^31^. We tested this gene in CHB+JPT populations. iSAFE rank, in 2Mbp around ADH1B gene, for the candidate mutation (rs1229984) is 8 (see Fig. S30B). The top rank mutation is an upstream mutation (rs3811801) 5kbp upstream of the candidate mutation rs1229984 and highly linked to it (*D*^*t*^ = 0.99). The second rank mutation (rs284787) is a 3^*t*^-UTR of ADH7 which is shown to be associated with Upper Aerodigestive Tract Cancers in a Japanese Population ^32^.

#### 6.4 Population specific selection: UK

The UK Biobank project was recently investigated for regions under selection. The regions were reported as a target of a recent selection by analyzing the structure of UK Biobank and Ancient Eurasians^27^. We applied iSAFE on GBR (British in England and Scotland) population in 1000GP to check if the favored mutation could be confirmed.

##### ATXN2-SH2B3

Galinsky et al. proposed a nonsynonymous mutation (rs3184504) as a candidate that is associated to blood pressure^33^. We tested this region in GBR population of 1000GP. This candidate mutation is jointly ranked first with two other mutations rs7137828, rs7310615 (see Fig. 3O, iSAFE = 0.27, *p*-val=1.6e-7). rs7137828 is an intronic mutation in ATXN2 that is associated with Primary Open Angle Glaucoma that is a leading cause of blindness worldwide ^34^. The other first rank mutation (rs7310615) is associated with blood expression levels of SH2B3^35^. Surprisingly, all of the top 10 mutations, ranked by iSAFE have a known association to a phenotype (Table S4), and are highly linked (Fig. S29).

##### CYP1A2/CSK

We tested a 5Mbp region around these genes in GBR population of 1000GP. The proposed mutation rs1378942 by^27^ with frequency 0.69 in GBR population is ranked 89 by iSAFE (iSAFE = 0.13, *p*-val=7.0e-5). The top-ranked mutation rs2470893 (Fig. 3U, iSAFE = 0.16, *p*-val=2.7e-5) is between CYP1A1 and CYP1A2 with frequency 0.40 in GBR and is associated with Caffeine metabolism ^11^. rs2470893 and rs1378942 are in a strong LD(*D*^*t*^ = 0.91).

##### FUT2

The signal of selection on 5Mbp around this region in GBR population is very weak (Fig. S30E), with peak iSAFE = 0.026, *p*-val=0.009. There is a very weak peak in 400kbp around FUT2 gene (chr:49077276-49475876). The stop gained mutation rs601338 proposed as a candidate mutation by^27^ is ranked 4 (*p*-val=0.1).

##### F12

The signal of selection on 5Mbp around this region in GBR population is very weak (Fig. S30F, peak iSAFE = 0.027, *p*-val=0.008). The proposed mutation rs2545801 has a very weak signal (*p*-val=0.2).

#### Other genes

##### PSCA

This gene has been reported as a target of selection in YRI population ^26^. A 5^*t*^UTR mutation rs2294008 proposed as a candidate favored mutation in this region that is associated with urinary bladder and gastric cancers ^36,37^. The signal of iSAFE in 5Mbp around this gene in YRI population is weak (see Fig. S30C, peak iSAFE = 0.04, *p*-val=2.4e-3). The proposed mutation rs2294008 is ranked 7 in 5Mbp region surrounding this region. The local rank in 400kbp around this gene is joint-first with 8 other mutations including rs2976392 which is also associated with diffuse-type gastric cancer ^37^. Other mutations are rs2978979, rs2920279, rs2978980, rs2920282, rs2294010, rs2717562, rs2978982. This 9 mutation are fully linked in YRI population in a 20kbp region that cover PSCA from upstream regulatory region to its down stream (chr8:143757286-143776668, GRCh37/hg19).

##### ASPM

This gene is reported to be a target of weak selection in GBR population^26^. The signal in 2Mbp around this gene is very weak (see Fig. S30D, peak-iSAFE = 0.025, *p*-val=0.01). The proposed mutation rs41310927 has a very weak signal (*p*-val=0.4). However, we do see a strong iSAFE signal 1.3Mbp away from the ASPM gene.

**Figure S23:**
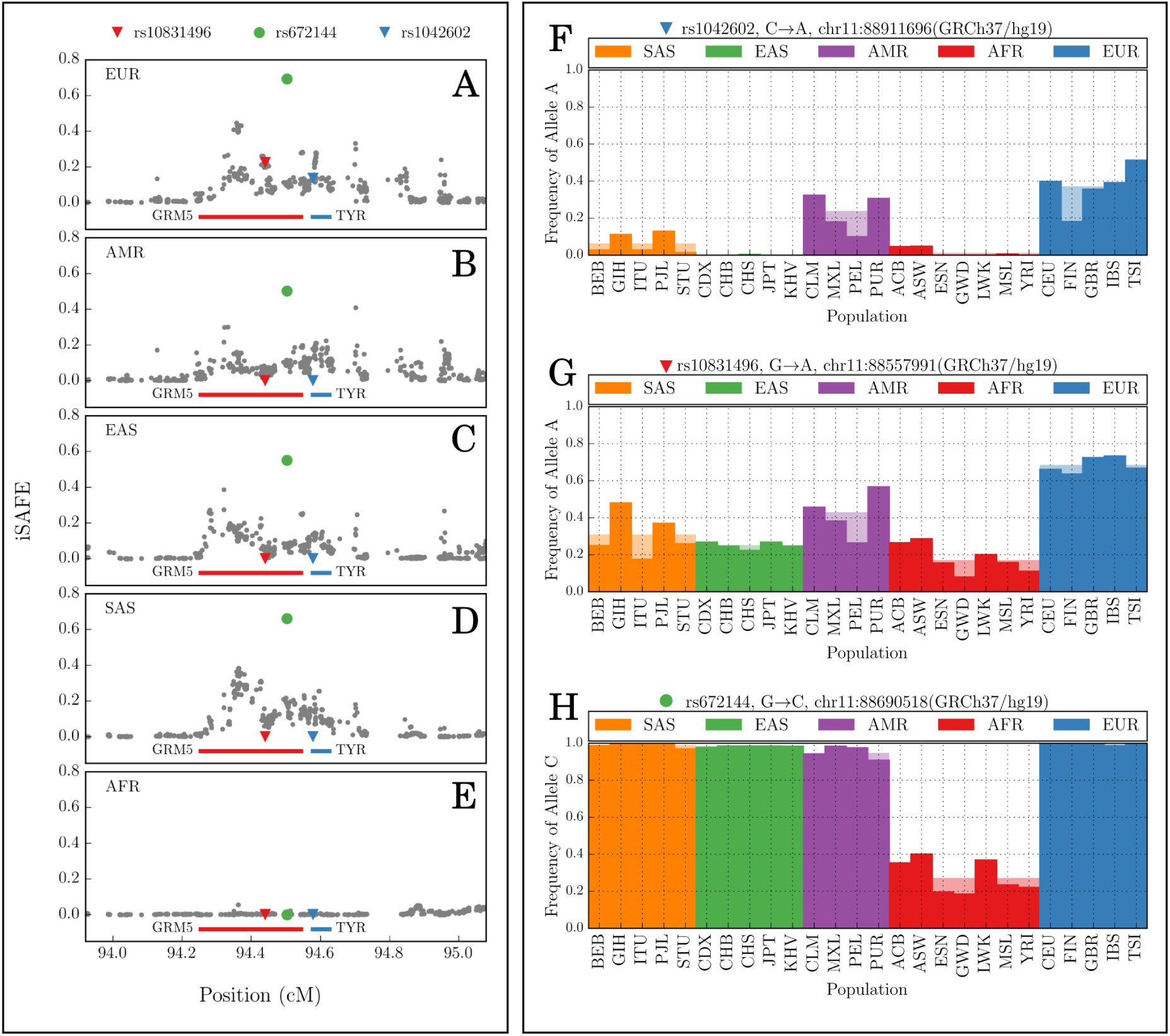
iSAFE on GRM5-TYR. The mutation rs672144 is the iSAFE top rank mutation in all the population of 1000GP except African that doesn’t show any signal of selection in this region.

**Figure S24:**
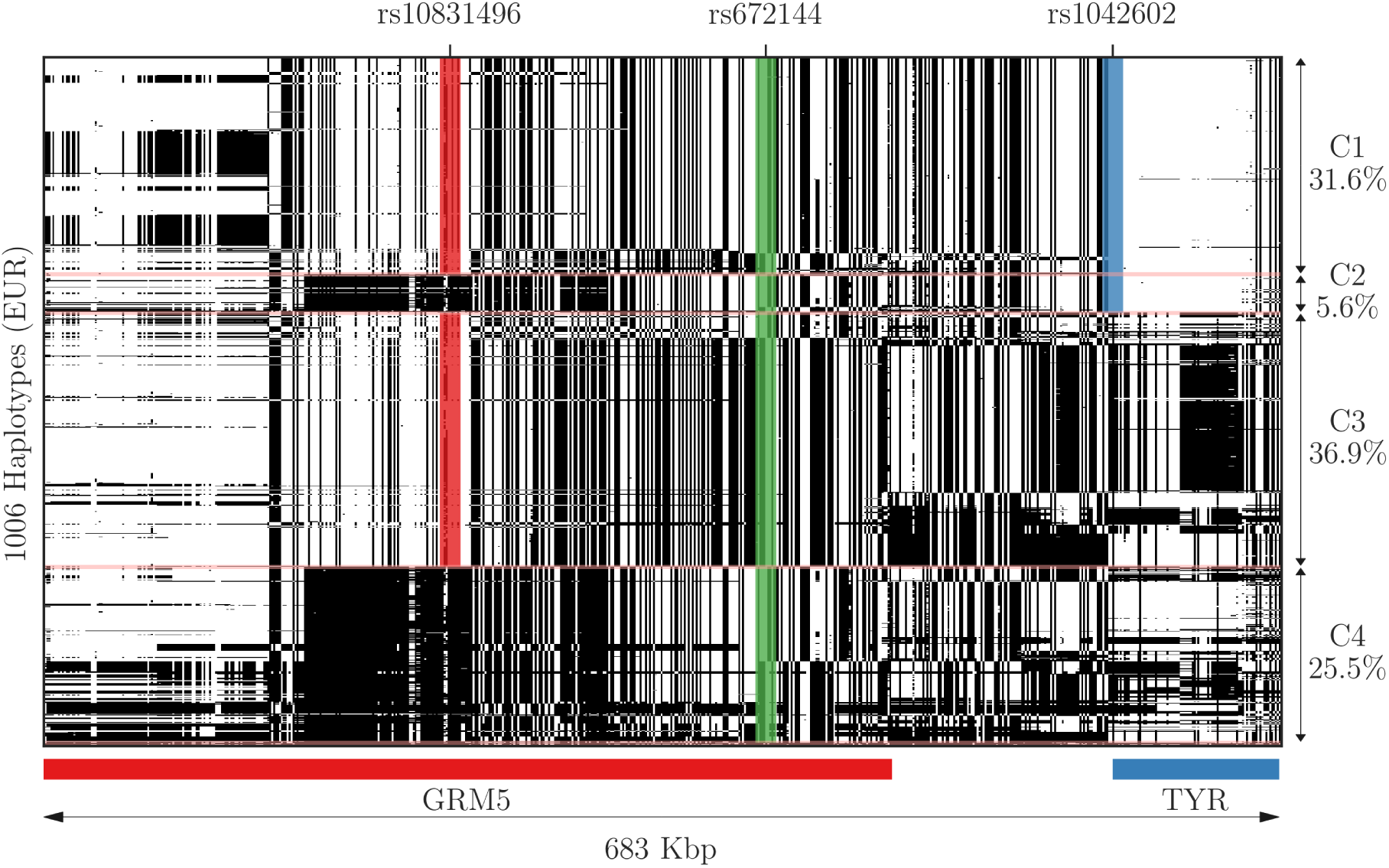
SNP matrix of GRM5-TYR. Each row is a haplotype and each column is a variant in EUR populations of 1000GP. In total we have 1006 haplotypes. Carriers haplotypes of derived alleles of rs10831496, rs672144, and rs1042602, are shaded by red, green, and blue, respectively. For making the plot sensible, We removed low frequency SNPs *f*_*EUR*_ < 0.2 and SNPs that are near fixation in the whole 1000GP, *f*_1000*GP*_ > 0.95.

**Figure S25:**
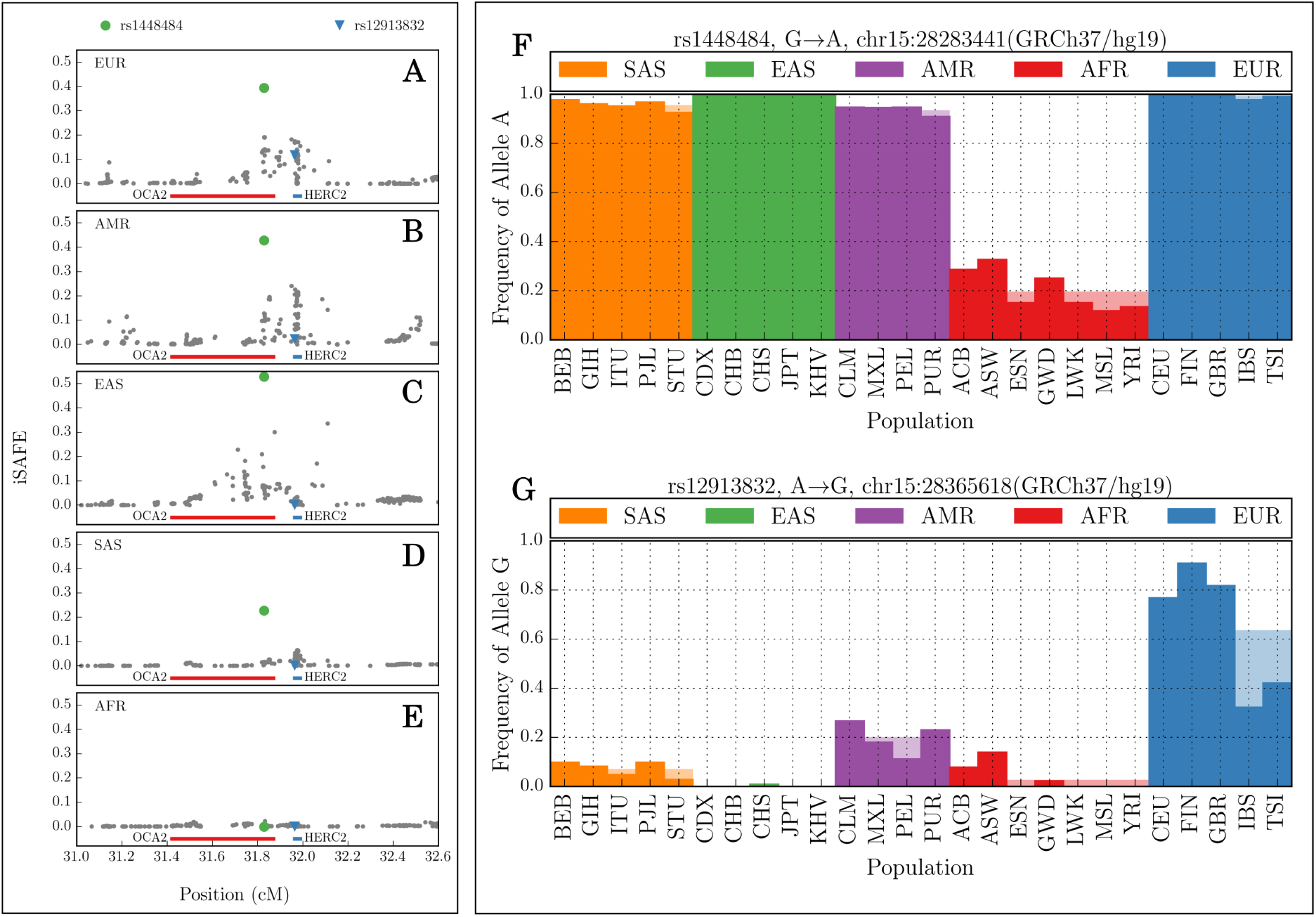
iSAFE on OCA2-HERC2. . The mutation rs1448484 is the iSAFE top rank mutation in all the population of 1000GP except African that doesn’t show any signal of selection in this region. rs12913832 is a candidate favored mutation for the selection in European, proposed by^9^. Table S2 provides iSAFE rank of some other candidate mutations associated with pigmentation^28^.

**Figure S26:**
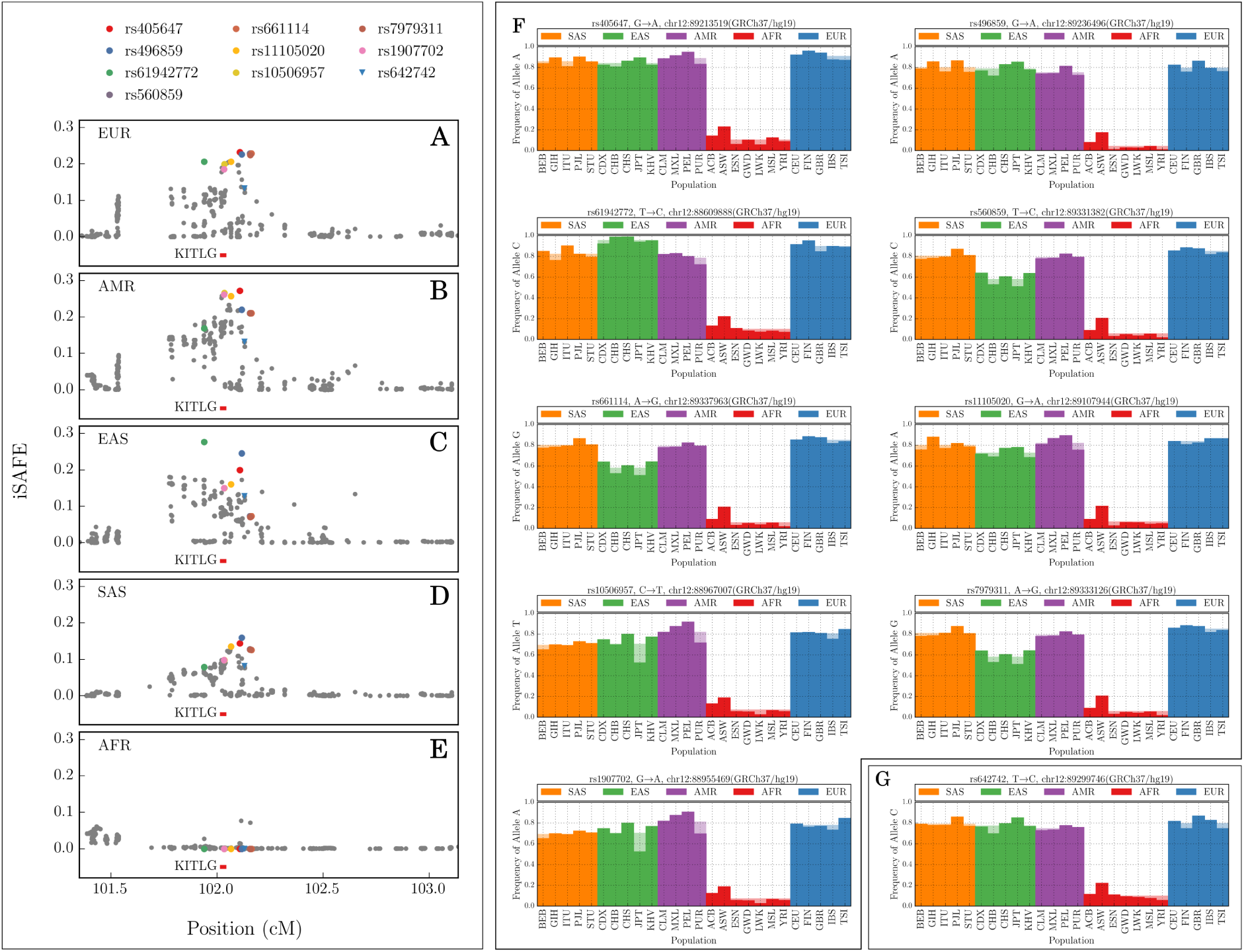
iSAFE on KITLG. iSAFE top rank mutations (circles) and candidate mutation rs642742 (blue triangle) proposed by^29^. See the Table S3 for more details.

**Figure S27:**
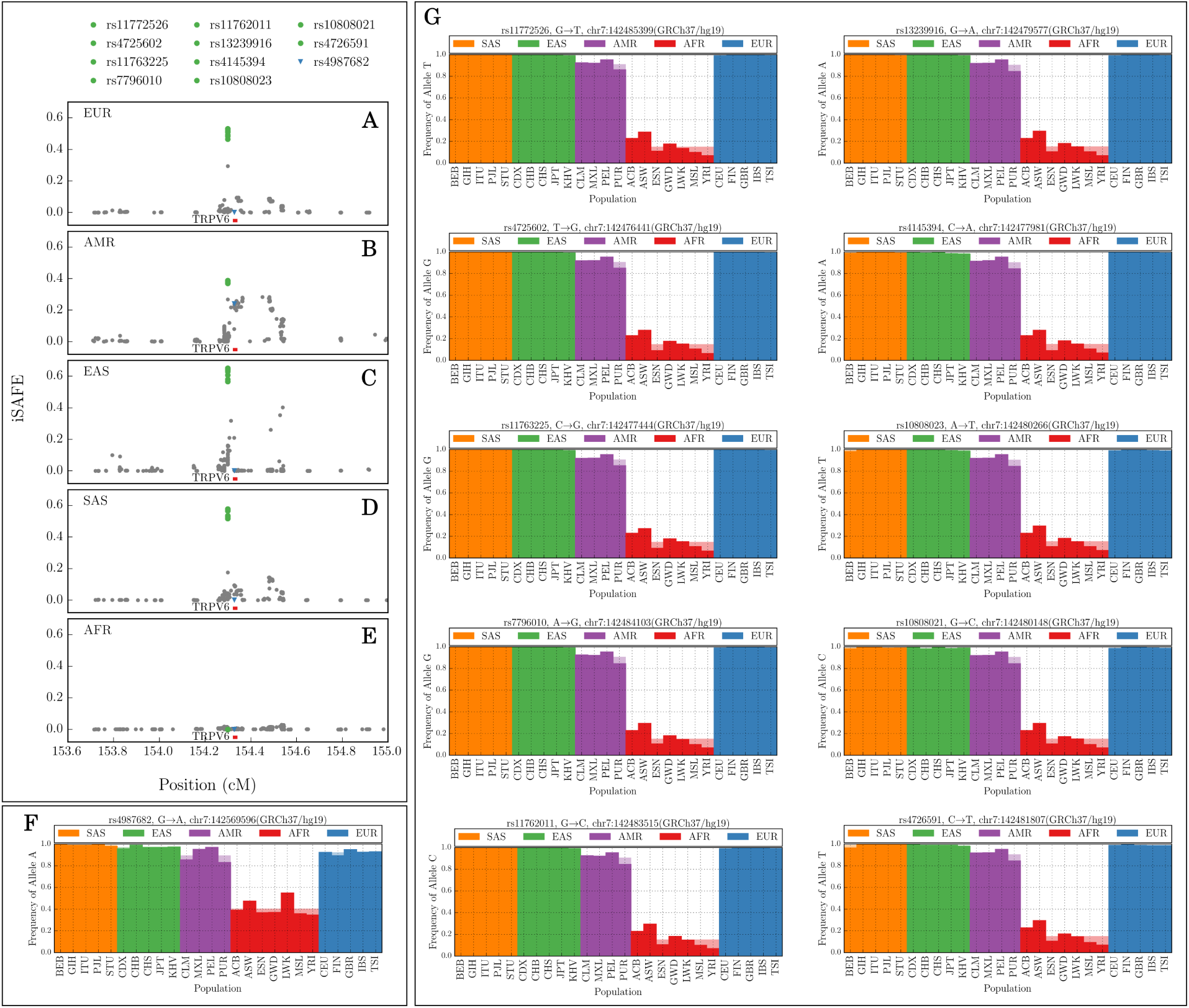
iSAFE on TRPV6. 10 mutations (rs11772526, rs4725602, rs11763225, rs7796010, rs11762011, rs13239916, rs4145394, rs10808023, rs10808021, and rs4726591) are highly linked and are top 10 iSAFE candidate mutations in all the 1000GP populations except for AFR where there is no signals of selection.

**Figure S28:**
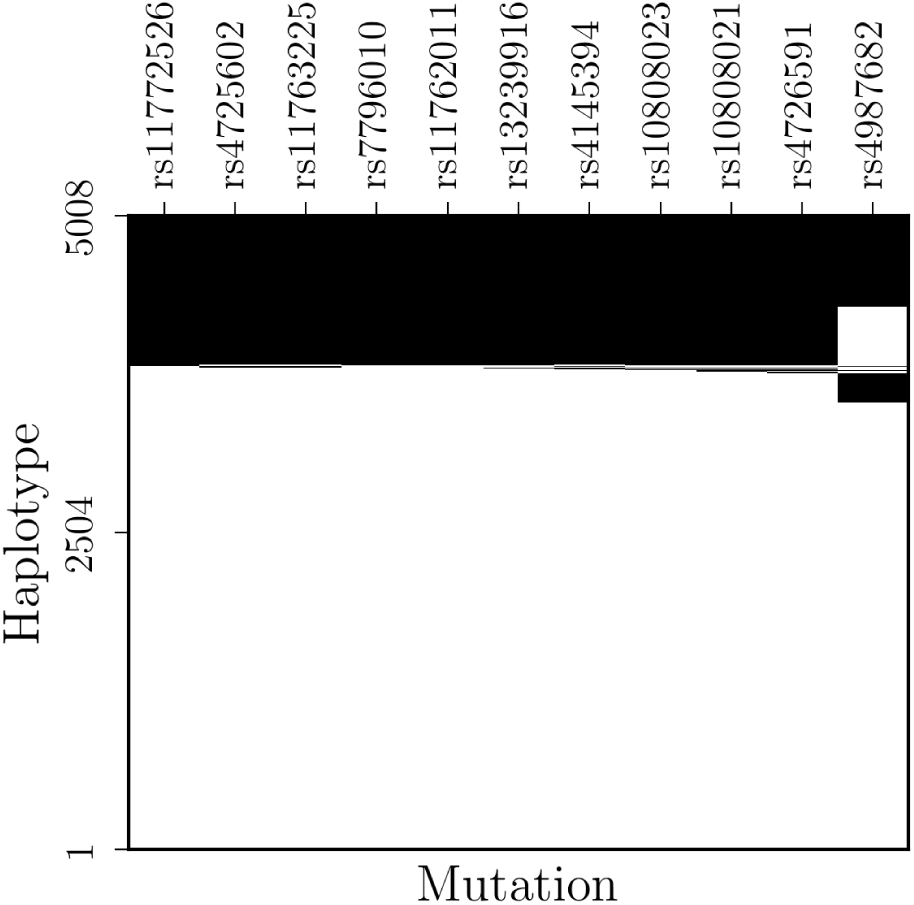
SNP matrix of TRPV6 top candidates. Haplotypes of top 10 iSAFE mutations, and the proposed mutation (rs4987682) by^26^, in 5Mbp around TRPV6 in 2504 *×* 2 haplotypes of 1000GP are shown. These mutations are sorted by their iSAFE rank from left to right. iSAFE top 10 mutations span a 9kbp region(chr7:142476441-142485399, GRCh37/hg19). White is derived and black is ancestral allele.

**Figure S29:**
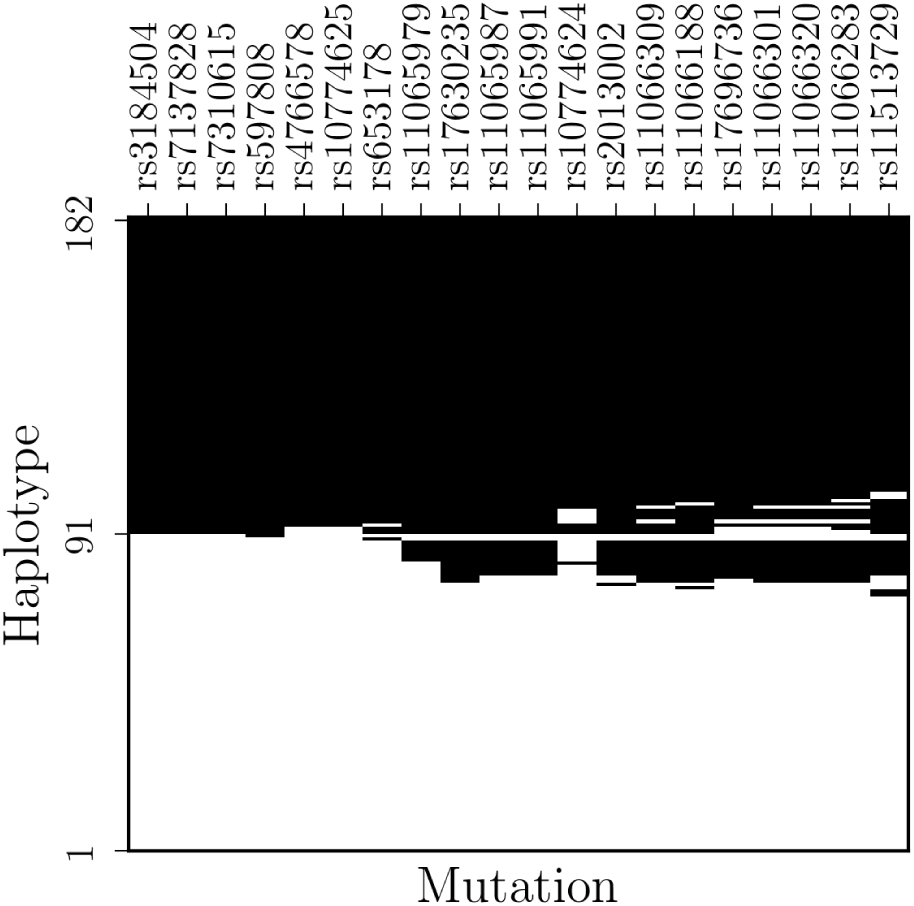
SNP matrix of ATXN2-SH2B3 top candidates. Haplotypes of top 20 iSAFE mutations in 5Mbp around ATXN2-SH2B3 in GBR population are shown. These mutations are sorted by their iSAFE rank from left to right. They span a 1.07Mbp region around ATXN2-SH2B3 region (chr12:111833788-112906415, GRCh37/hg19). White is derived and black is ancestral allele. Most of these mutations are associated to a phenotype (see Table S4).

**Figure S30:**
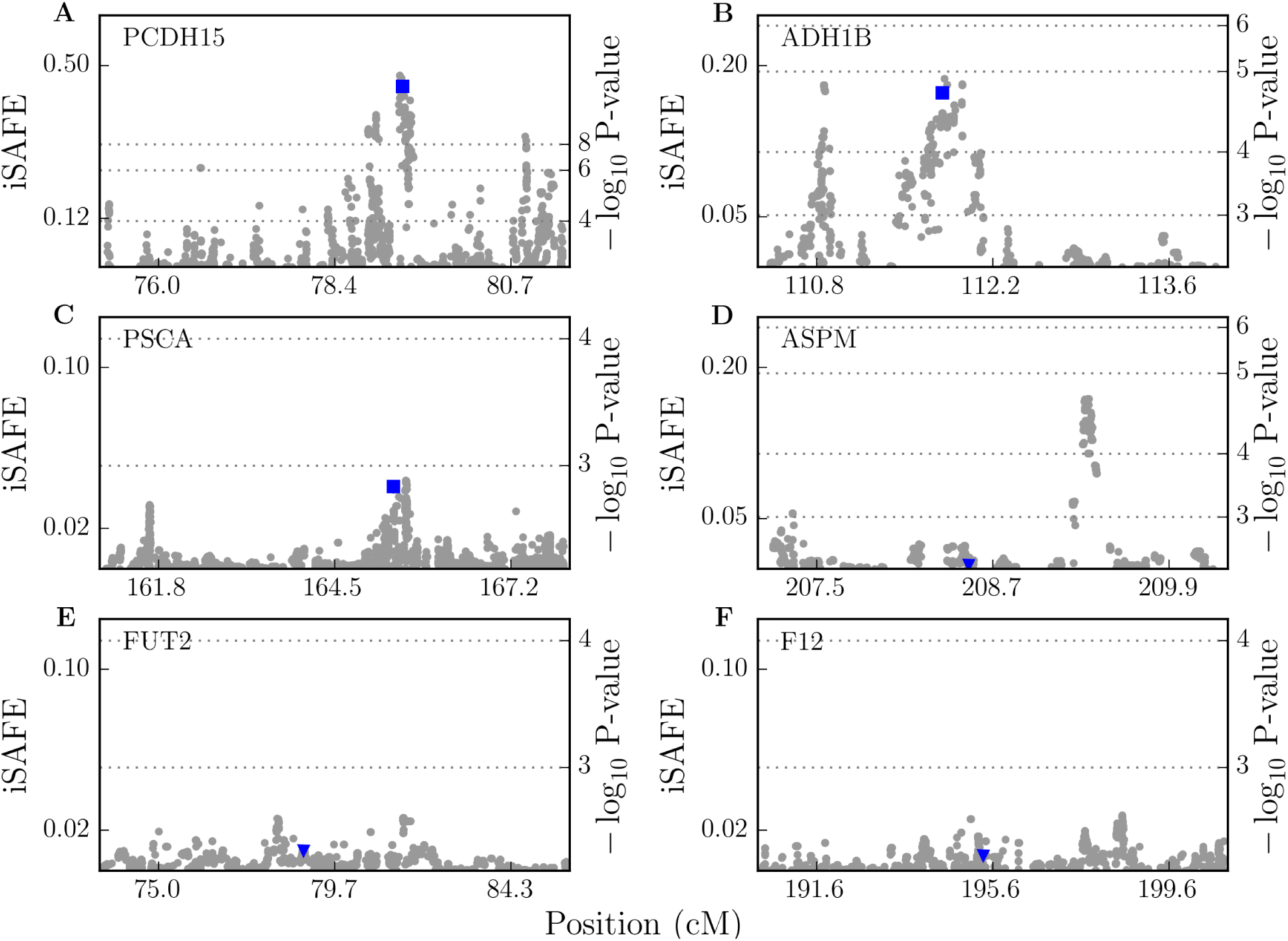
iSAFE on Targets of Selection. iSAFE-scores on regions under selection. Putative favored mutation is shown in blue square when it is among iSAFE top rank mutations, and in blue triangle when the signal of selection is very weak.

**Table S1:** iSAFE on 8 well characterized selective sweeps.

| Gene | Target Population | Candidate SNP ID | Candidate SNP Function | Frequency | Selective Advantage | iSAFE Rank | P-value | Selection Reference | Functional Reference |
| --- | --- | --- | --- | --- | --- | --- | --- | --- | --- |
| SLC24A5 | CEU | rs1426654 | Missense | 1 | Light skin pigmentation | 1 | <1.3e-8 | <sup>4</sup> | <sup>38</sup> |
| EDAR | CHB+JPT | rs3827760 | Missense | 0.87 | Hair and teeth | 1 | <1.3e-8 | <sup>4</sup> | <sup>39,40</sup> |
| LCT/MCM6 | FIN | rs4988235 | Intron | 0.59 | Lactase persistence | 1 | <1.3e-8 | <sup>26</sup> | <sup>21,41</sup> |
| TLR1 | CEU | rs5743618 | Missense | 0.77 | Sepsis, leprosy, tuberculosis | 1 | 1.0e-5 | <sup>24</sup> | <sup>42</sup> |
| ACKR1/DARC | YRI | rs2814778 | 5'UTR | 1 | Malaria resistance | 1 | 2.8e-5 | <sup>43</sup> | <sup>44</sup> |
| ABCC11 | CHB+JPT | rs17822931 | Missense | 0.93 | Cold climate, earwax, body odour | 2 | <1.3e-8 | <sup>22</sup> | <sup>22</sup> |
| HBB | YRI | rs334 | Missense | 0.14 | Malaria resistance | 4 | 1.6e-4 | <sup>20</sup> | <sup>45</sup> |
| G6PD | YRI | rs1050828 | Missense | 0.21 | Malaria resistance | 13 | 7.3e-6 | <sup>26</sup> | <sup>23</sup> |

**Table S2:** iSAFE rank of putative favored variants of OCA2-HERC2. iSAFE rank of candidate mutations proposed by ^9,28^ in 1Mbp region around OCA2-HERC2 that are associated with eye, hair, and skin pigmentation.

| ID | Association | Population | iSAFE Rank | P-Value |
| --- | --- | --- | --- | --- |
| rs916977 | Blue eye | CEU | 15 | 4.1E-5 |
| rs1667394 | Blue eye & blond hair | CEU | 16 | 4.3E-5 |
| rs1129038 | Blue eye | CEU | 21 | 6.2E-5 |
| rs12913832 | Blue eye, skin & hair | CEU | 21 | 6.2E-5 |
| rs4778138 | Blue eye | CEU | 70 | 1.6E-4 |
| rs4778241 | Blue eye | CEU | 72 | 1.8E-4 |
| rs1800414 | Skin | CHB+JPT | 122 | 2.6E-3 |

**Table S3:** KITLG candidate variants. iSAFE rank of top mutations in 2 Mbp around KITLG gene. sorted by their Mean Reciprocal Ranks, calculated over EUR, SAS, EAS, AMR. Only those with Mean Reciprocal Rank greater than 0.1 are shown (the candidate mutation rs642742 proposed by^29^ is also reported in the last row). Frequency and iSAFE score for this region in all the 1000GP populations are provided in S26.

| ID | iSAFE Rank<br>EUR | iSAFE Rank<br>SAS | iSAFE Rank<br>EAS | iSAFE Rank<br>AMR | Mean Reciprocal Rank<br>EUR, SAS, EAS, AMR | iSAFE Rank<br>CEU |
| --- | --- | --- | --- | --- | --- | --- |
| rs405647 | 1 | 2 | 3 | 1 | 0.71 | 1 |
| rs496859 | 4 | 1 | 2 | 12 | 0.46 | 7 |
| rs61942772 | 10 | 57 | 1 | 94 | 0.28 | 22 |
| rs560859 | 2 | 4 | 152 | 20 | 0.2 | 5 |
| rs661114 | 2 | 6 | 151 | 20 | 0.18 | 5 |
| rs11105020 | 8 | 3 | 32 | 5 | 0.17 | 23 |
| rs10506957 | 17 | 22 | 46 | 2 | 0.16 | 2 |
| rs7979311 | 5 | 5 | 156 | 20 | 0.11 | 3 |
| rs1907702 | 22 | 20 | 45 | 3 | 0.11 | 8 |
| rs642742 | 30 | 49 | 64 | 166 | 0.02 | 94 |

**Table S4:** ATXN2-SH2B3 candidate variants. iSAFE rank of top 20 mutations in GBR population of 1000GP in 5Mbp around ATXN2-SH2B3 region and their association to diseases.

| ID | Rank | P-value | Gene | Function | GBR Frequency | Association | Reference |
| --- | --- | --- | --- | --- | --- | --- | --- |
| rs3184504 | 1 | 2.2e-7 | SH2B3 | missense | 0.5 | Blood pressure and hypertension, Coronary artery disease, & more | <sup>46</sup> |
| rs7137828 | 1 | 2.2e-7 | ATXN2 | intron | 0.5 | Primary open-angle glaucoma | <sup>34</sup> |
| rs7310615 | 1 | 2.2e-7 | SH2B3 | intron | 0.5 | Fibrinogen levels | <sup>35</sup> |
| rs597808 | 4 | 2.7e-7 | ATXN2 | intron | 0.49 | Systemic lupus erythematosus | <sup>47</sup> |
| rs4766578 | 5 | 3.0e-7 | ATXN2 | intron | 0.51 | Vitiligo | <sup>48</sup> |
| rs10774625 | 5 | 3.0e-7 | ATXN2 | intron | 0.51 | Systemic lupus erythematosus, Retinal vascular caliber | <sup>47</sup> |
| rs653178 | 7 | 3.1e-7 |  | regulatory | 0.5 | Blood pressure and hypertension, Myocardial infarction, & more | <sup>46</sup> |
| rs11065979 | 8 | 4.4e-7 |  | intergenic | 0.47 | Cancer (pleiotropy) | <sup>49</sup> |
| rs17630235 | 9 | 4.6e-7 | TRAFD1 | downstream | 0.43 | Body mass index | <sup>50</sup> |
| rs11065987 | 10 | 4.9e-7 |  | intergenic | 0.45 | Tetralogy of Fallot, Coronary artery disease, & more | <sup>51</sup> |
| rs11065991 | 10 | 4.9e-7 | BRAP | intron | 0.45 |  |  |
| rs10774624 | 12 | 5.2e-7 | RP3-473L9.4 | intron,nc | 0.52 | Rheumatoid arthritis | <sup>52</sup> |
| rs2013002 | 13 | 8.2e-7 | ALDH2 | upstream | 0.44 |  |  |
| rs11066309 | 14 | 1.1e-6 | PTPN11 | intron | 0.45 |  |  |
| rs11066188 | 15 | 1.5e-6 |  |  | 0.43 |  |  |
| rs17696736 | 16 | 1.5e-6 | NAA25 | intron | 0.46 | Ischemic stroke, Type 1 diabetes, & more | <sup>53</sup> |
| rs11066301 | 17 | 1.9e-6 | PTPN11 | intron | 0.46 | Hematological parameters | <sup>54</sup> |
| rs11066320 | 17 | 1.9e-6 | PTPN11 | intron | 0.46 |  |  |
| rs11066283 | 19 | 2.1e-6 | RPL6 | downstream | 0.46 |  |  |
| rs11513729 | 20 | 2.2e-6 | MAPKAPK5-AS1 | downstream | 0.45 |  |  |

**Table S5:** TRPV6 candidate variants. iSAFE rank of top mutations in 5Mbp around TRPV6 gene. sorted by their Mean Reciprocal Ranks, calculated over EUR, SAS, EAS, AMR.

| ID | iSAFE Rank<br>EUR | iSAFE Rank<br>SAS | iSAFE Rank<br>EAS | iSAFE Rank<br>AMR | Mean Reciprocal Rank<br>EUR, SAS, EAS, AMR | iSAFE Rank<br>CEU |
| --- | --- | --- | --- | --- | --- | --- |
| rs11772526 | 4.0 | 1.0 | 1.0 | 1.0 | 0.81 | 4.0 |
| rs4725602 | 1.0 | 4.0 | 1.0 | 2.0 | 0.69 | 1.0 |
| rs11763225 | 1.0 | 4.0 | 5.0 | 2.0 | 0.49 | 1.0 |
| rs7796010 | 4.0 | 1.0 | 3.0 | 6.0 | 0.44 | 4.0 |
| rs11762011 | 4.0 | 3.0 | 3.0 | 6.0 | 0.27 | 4.0 |
| rs13239916 | 4.0 | 6.0 | 6.0 | 4.0 | 0.21 | 4.0 |
| rs4145394 | 3.0 | 8.0 | 10.0 | 5.0 | 0.19 | 1.0 |
| rs10808023 | 8.0 | 7.0 | 7.0 | 8.0 | 0.13 | 4.0 |
| rs10808021 | 9.0 | 10.0 | 8.0 | 8.0 | 0.12 | 10.0 |
| rs4726591 | 10.0 | 9.0 | 9.0 | 10.0 | 0.11 | 4.0 |

## References

[1] Vitti, J. J., Grossman, S. R. & Sabeti, P. C. Detecting natural selection in genomic data. Annual review of genetics 47, 97–120 (2013).

[2] Fan, S., Hansen, M. E., Lo, Y. & Tishko, S. A. Going global by adapting local: A review of recent human adaptation. Science 354, 54–59 (2016).

[3] Field, Y. et al. Detection of human adaptation during the past 2000 years. Science 354 60–764 (2016).

[4] Grossman, S. R. et al. A composite of multiple signals distinguishes causal variants in regionsof positive selection. Science 327, 883–886 (2010).

[5] Schrider, D. R., Mendes, F. K., Hahn, M. W. & Kern, A. D. Soft shoulders ahead: spurious signatures of soft and partial selective sweeps result from linked hard sweeps. Genetics 200, 267–284 (2015).

[6] Ronen, R. et al. Predicting carriers of ongoing selective sweeps without knowledge of the-favored allele. PLoS Genet 11, e1005527 (2015).

[7] Wang, M. et al. Detecting recent positive selection with high accuracy and reliability by-conditional coalescent tree. Molecular biology and evolution msu 244 (2014)

[8] Voight, B. F., Kudaravalli, S., Wen, X. & Pritchard, J. K. A map of recent positive selectionin the human genome. PLoS Biol 4, e72 (2006).

[9] Wilde, S. et al. Direct evidence for positive selection of skin, hair, and eye pigmentation inEuropeans during the last 5,000 y. Proceedings of the National Academy of Sciences 111,4832–4837 (2014).

[10] Beleza, S. et al. Genetic architecture of skin and eye color in an African-European admixedpopulation. PLoS Genet 9, e1003372 (2013).

[11] Cornelis, M. C. et al. Genome-wide meta-analysis identi_es six novel loci associated withhabitual co_ee consumption. Molecular psychiatry 20, 647-656 (2015).-

[12] Fu, Y.-X. Statistical properties of segregating sites. Theoretical population biology 48, 172–197(1995).

[13] Wiuf, C. & Donnelly, P. Conditional genealogies and the age of a neutral mutant. Theoretical population biology 56, 183–201 (1999).

[14] Ewing, G. & Hermisson, J. MSMS: a coalescent simulation program including recombination, demographic structure and selection at a single locus. Bioinformatics 26, 2064–2065 (2010).

[15] Nachman, M. W. & Crowell, S. L. Estimate of the mutation rate per nucleotide in humans. Genetics 156, 297–304 (2000).

[16] Campbell, C. D. et al. Estimating the human mutation rate using autozygosity in a founder population. Nature genetics 44, 1277–1281 (2012).

[17] Jensen-Seaman, M. I. et al. Comparative recombination rates in the rat, mouse, and human genomes. Genome research 14, 528–538 (2004).

[18] Gravel, S. et al. Demographic history and rare allele sharing among human populations. Proceedings of the National Academy of Sciences 108, 11983–11988 (2011)

[19] Szpiech, Z. A. & Hernandez, R. D. selscan: an e_cient multithreaded program to perform EHH-based scans for positive selection. Molecular biology and evolution 31, 2824–2827 (2014).

[20] Sabeti, P. C. et al. Positive natural selection in the human lineage. Science 312, 1614–1620(2006).

[21] Enattah, N. S. et al. Identification of a variant associated with adult-type hypolactasia. Nature genetics 30, 233–237 (2002).

[22] Ohashi, J., Naka, I. & Tsuchiya, N. The impact of natural selection on an ABCC11 SNP determining earwax type. Molecular biology and evolution 28, 849–857 (2011).

[23] Tishko, S. A. et al. Haplotype diversity and linkage disequilibrium at human G6PD: recentorigin of alleles that confer malarial resistance. Science 293, 455–462 (2001).

[24] Heelnger, C. et al. Haplotype structure and positive selection at TLR1. European Journal of Human Genetics 22, 551–557 (2014).

[25] Coop, G. et al. The role of geography in human adaptation. PLoS Genet 5, e1000500 (2009).

[26] Peter, B. M., Huerta-Sanchez, E. & Nielsen, R. Distinguishing between selective sweeps fromstanding variation and from a de novo mutation. PLoS Genet 8, e1003011 (2012).

[27] Galinsky, K. J., Loh, P.-R., Mallick, S., Patterson, N. J. & Price, A. L. Population structureof UK Biobank and ancient Eurasians reveals adaptation at genes inuencing blood pressure. The American Journal of Human Genetics 99, 1130–1139 (2016).

[28] Donnelly, M. P. et al. A global view of the OCA2-HERC2 region and pigmentation. Human genetics 131, 683–696 (2012).

[29] Miller, C. T. et al. cis-Regulatory changes in Kit ligand expression and parallel evolution ofpigmentation in sticklebacks and humans. Cell 131, 1179–1189 (2007).

[30] Suzuki, Y. et al. Gain-of-function haplotype in the epithelial calcium channel TRPV6 is a riskfactor for renal calcium stone formation. Human molecular genetics 17, 1613–1618 (2008).

[31] Park, B. L. et al. Extended genetic e_ects of ADH cluster genes on the risk of alcohol dependence: from GWAS to replication. Human genetics 132, 657–668 (2013).

[32] Oze, I. et al. Impact of multiple alcohol dehydrogenase gene polymorphisms on risk of upperaerodigestive tract cancers in a Japanese population. Cancer Epidemiology and Prevention Biomarkers 18, 3097–3102 (2009).

[33] for Blood Pressure Genome-Wide Association Studies, I. C. et al. Genetic variants in novelpathways inuence blood pressure and cardiovascular disease risk. Nature 478, 103–109 (2011).

[34] Bailey, J. N. C. et al. Genome-wide association analysis identi_es TXNRD2, ATXN2 andFOXC1 as susceptibility loci for primary open-angle glaucoma. Nature genetics (2016).

[35] De Vries, P. S. et al. A meta-analysis of 120,246 individuals identi_es 18 new loci for brinogen concentration. Human molecular genetics ddv454 (2015).

[36] Wu, X. et al. Genetic variation in the prostate stem cell antigen gene PSCA confers susceptibility to urinary bladder cancer. Nature genetics 41, 991–995 (2009).

[37] Sakamoto, H. et al. Genetic variation in PSCA is associated with susceptibility to di_usetype gastric cancer. Nature genetics 40, 730–740 (2008).

[38] Sturm, R. A. & Duy, D. L. Human pigmentation genes under environmental selection. Genome biology 13, 248 (2012).

[39] Fujimoto, A. et al. A replication study con_rmed the EDAR gene to be a major contributor to population di_erentiation regarding head hair thickness in Asia. Human genetics 124, 179–185(2008).

[40] Bryk, J. et al. Positive selection in East Asians for an EDAR allele that enhances NF-B activation. PLoS One 3, e2209 (2008).

[41] Olds, L. C. & Sibley, E. Lactase persistence DNA variant enhances lactase promoter activity in vitro: functional role as a cis regulatory element. Human molecular genetics 12, 2333–2340 (2003).

[42] Wong, S. H. et al. Leprosy and the adaptation of human toll-like receptor 1. PLoS Pathog 6,e1000979 (2010).

[43] McManus, K. F. et al. Population genetic analysis of the DARC locus (Duy) reveals adaptation from standing variation associated with malaria resistance in humans. PLoS Genetics 13, e1006560 (2017).

[44] Miller, L. H., Mason, S. J., Clyde, D. F. & McGinniss, M. H. The resistance factor to Plasmodium vivax in blacks: the Duy-blood-group genotype, FyFy. New England Journal of Medicine 295, 302–304 (1976).

[45] Network, M. G. E. Reappraisal of known malaria resistance loci in a large multicenter study. Nature genetics 46, 1197–1204 (2014).

[46] Levy, D. et al. Genome-wide association study of blood pressure and hypertension. Nature genetics 41, 677–687 (2009)

[47] Bentham, J. et al. Genetic association analyses implicate aberrant regulation of innate and adaptive immunity genes in the pathogenesis of systemic lupus erythematosus. Nature genetics(2015).

[48] Jin, Y. et al. Genome-wide association analyses identify 13 new susceptibility loci for generalized vitiligo. Nature genetics 44, 676–680 (2012).

[49] Fehringer, G. et al. Cross-CancerGenome-Wide Analysis of Lung, Ovary, Breast, Prostate, and Colorectal Cancer Reveals Novel Pleiotropic Associations. Cancer research 76, 5103-5114(2016).

[50] Locke, A. E. et al. Genetic studies of body mass index yield new insights for obesity biology. Nature 518, 197–206 (2015).

[51] Cordell, H. J. et al. Genome-wide association study identi_es loci on 12q24 and 13q32 associated with tetralogy of Fallot. Human molecular genetics dds552 (2013).

[52] Okada, Y. et al. Genetics of rheumatoid arthritis contributes to biology and drug discovery. Nature 506, 376–381 (2014).

[53] Dichgans, M. et al. Shared genetic susceptibility to ischemic stroke and coronary artery disease. Stroke 45, 24–36 (2014).

[54] Soranzo, N. et al. A genome-wide meta-analysis identi_es 22 loci associated with eight hematological parameters in the HaemGen consortium. Nature genetics 41, 1182–1190 (2009).

